# Evaluating the role of coevolution in a horizontally transmitted mutualism

**DOI:** 10.1101/2021.12.04.471243

**Authors:** Kayla Stoy, Joselyne Chavez, Valeria De Las Casas, Venkat Talla, Aileen Berasategui, Levi Morran, Nicole Gerardo

## Abstract

Mutualism depends on the alignment of host and symbiont fitness. Horizontal transmission can readily decouple fitness interests, yet horizontally transmitted mutualisms are common in nature. We hypothesized that pairwise coevolution and specialization in host-symbiont interactions underlies the maintenance of cooperation in a horizontally transmitted mutualism. Alternatively, we predicted selection by multiple host species may select for cooperative traits in a generalist symbiont through diffuse coevolution. We tested for signatures of pairwise coevolutionary change between the squash bug *Anasa tristis* and its horizontally acquired bacterial symbiont *Caballeronia spp*. by measuring local adaptation. We found no evidence for local adaptation between sympatric combinations of *A. tristis* squash bugs and *Caballeronia spp.* across their native geographic range. To test for diffuse coevolution, we performed reciprocal inoculations to test for specialization between three *Anasa* host species and *Caballeronia spp.* symbionts isolated from conspecific hosts. We observed no evidence of specialization across host species. Our results demonstrate generalist dynamics underlie the interaction between *Anasa* insect hosts and their *Caballeronia spp.* symbionts. Specifically, diffuse coevolution between multiple host species with a shared generalist symbiont may maintain cooperative traits despite horizontal transmission.

## Introduction

Hosts across all domains of life form essential mutualistic interactions with microbial symbionts. Mutualistic microbes fulfill a variety of integral functions for host development and survival, including augmenting nutrition, providing defense against pathogens, and facilitating development (Pais et al. 2008; Boucias et al. 2012; Salem et al. 2013; Masson et al. 2015; Vorburger and Perlman 2018; Gerardo et al. 2020; Kaltenpoth and Flórez 2020). Likewise, symbionts may depend on hosts for nutrition, replication without competition, and transmission (Lee & Ruby, 1994; Prell *et al.*, 2009; Macdonald *et al.*, 2012; Wollenberg & Ruby, 2012; but see also Garcia & Gerardo, 2014). The persistence of cooperative interactions relies on the alignment of host and symbiont fitness. Vertical transmission facilitates fitness alignment because symbiont transmission depends on host survival and reproduction (Bull 1994; Herre et al. 1999; Shapiro and Turner 2018). In contrast, horizontal transmission can misalign host and symbiont fitness. Horizontally transmitted symbionts can survive outside their hosts, relaxing their dependence on host fitness and potentially increasing selection for transmission that increases within host replication and exploitation (Anderson and May 1982; Ewald 1987; Bull 1994; Frank 1994; Wade 2007). Furthermore, horizontal transmission may leave mutualisms vulnerable to breakdown if selection from the environment favors traits for survival apart from mutualistic partners or environmental change makes symbionts difficult to acquire (Sachs and Simms 2006). Despite these potential costs, horizontal transmission is common across symbiotic mutualisms (Lee and Ruby 1994*a*; Wilkinson 1997; Simms 2002; Mikheyev et al. 2006; Kikuchi et al. 2011; Henry et al. 2013; Chrostek et al. 2017; Hartmann et al. 2017; Li et al. 2017). How do horizontally transmitted mutualisms persist given these opportunities for costs and evolutionary breakdown?

Coevolution underlies both antagonistic and mutualistic interspecific interactions (Dybdahl and Lively 1998; Moran 2001; Currie et al. 2003; Decaestecker et al. 2007; Janzen 2014; Wilson and Duncan 2015). Vertical transmission increases the likelihood for pairwise coevolution because interactions between specific host and symbiont lineages are conserved through time. These conditions can promote fitness alignment by preserving opportunities for reciprocal selection between host and symbiont lineages across generations. Reciprocal genomic and phenotypic changes underlying transmission and fitness are often observed across vertically transmitted mutualisms, demonstrating evidence for coevolution (Riegler et al. 2004; Toft and Andersson 2010; Ilinsky 2013; Wilson and Duncan 2015; Lee et al. 2020). Opportunities for coevolution, however, are not limited to vertically transmitted interactions. Coevolution is also presumed to occur frequently in horizontally transmitted mutualisms due to the prevalence of dependency, genetic specificity, and phylogenetic co-diversification between hosts and horizontally transmitted symbionts (Lee and Ruby 1994*b*; Aanen et al. 2002; Garcia-Cuetos et al. 2005; Brucker and Bordenstein 2012; Wang et al. 2012; Murfin et al. 2015; Parker et al. 2017; Forsman et al. 2020). Furthermore, substantial empirical evidence demonstrates an important role for coevolution across horizontally transmitted host-parasite symbioses (Kaltz et al. 1999; Fischer and Foitzik 2004; Lively et al. 2004; Decaestecker et al. 2007; Greischar and Koskella 2007; King et al. 2009; Morran et al. 2011; Thrall et al. 2012; Luijckx et al. 2013). Empirical evidence suggests coevolution can even result in reduced antagonism within symbiotic interactions, including under horizontal transmission (Gibson et al. 2015; Nelson and May 2017). Therefore, we hypothesize that coevolution may also underlie the long-term maintenance of horizontally transmitted mutualisms.

Horizontal transmission often presents greater inherent risks for hosts than their microbial symbionts. Hosts that depend on microbial symbionts but fail to acquire symbionts or acquire exploitative symbiont genotypes exhibit reduced fitness (Bidartondo and Bruns 2001; Kikuchi et al. 2009; Sachs et al. 2010; Boucias et al. 2012). However, horizontally transmitted mutualisms are common. This prevalence and long-term persistence of these interactions suggests pathways exists by which hosts and their horizontally transmitted symbionts align their fitness interests. Specialization through tight pairwise coevolution presents one potential pathway. Spatial structure may promote pairwise coevolution by conserving interactions between specific host and symbiont lineages across generations (Wilkinson 2001). Hosts that exert strong selection on their local microbial symbionts for cooperative traits may minimize exploitation. In turn, symbionts that reciprocate selection on local host lineages to provide fitness benefits may evolve greater reliance on these hosts, increasing their investment in the interaction. Specialization can align partner fitness by increasing the rewards available through the interaction (Douglas 1998; Schwartz and Hoeksema 1998). As a result, pairwise coevolution that produces specialized interactions across spatially structured populations may reduce the costs of horizontal transmission by facilitating fitness alignment.

Despite its potential benefits, specialization may also limit the pool of potential partners with which hosts can interact. Specialization may then result in an evolutionary dead end for hosts if the environment changes and decreases the prevalence of compatible symbiont genotypes (Futuyma and Moreno 1988; Douglas 1998; Sachs and Simms 2006). Meanwhile, hosts that interact with a range of generalist symbionts are more likely to find compatible partners. Moreover, long-term interactions between hosts and generalist symbionts may promote diffuse coevolution, a potential pathway by which the costs of the horizontal transmission of generalist symbionts may be limited. Diffuse coevolution results through reciprocal selection between a range of mutualistic partners with their shared common partner (Hougen-Eitzman and Rausher 1994; Iwao and Rausher 1997). In mutualism, for example, diffuse coevolution may occur through selection between a common generalist symbiont with a range of host species. Through diffuse coevolutionary interactions, the exchange of benefits is maintained, and exploitation is limited, via reciprocal selection for cooperative traits between the symbiont and its range of hosts. Diffuse coevolution may align fitness interests because symbionts interacting with a range of hosts are under constant selection for cooperative traits, and selection from multiple hosts limits opportunities for symbiont to become highly adapted to exploit any single host. Symbionts may also benefit from diffuse coevolution with a range of hosts by increasing their own transmission potential while avoiding exploitation and sequestration by a host. As a result, diffuse coevolution between hosts and generalist symbionts has potential to align host and symbiont fitness and limit the costs of horizontal transmission, especially in variable or unstable environments where specificity can reduce the probability of finding a compatible partner.

Patterns of host-symbiont specialization (or lack thereof) can provide evidence of whether pairwise or diffuse coevolution contribute to mutualistic interactions. Host-symbiont specialization is a signature of pairwise coevolution (Thompson 1994). Pairwise coevolution promotes specialization because hosts and symbionts achieve the greatest fitness advantages by imposing constant strong selection on specific partners. Specialization between hosts and symbionts can be observed across different levels of host and symbiont interactions, including within and across species. For mutualistic interactions, specialization results when specific host and symbiont combinations produce higher fitness interactions than those attained through alternative host-symbiont combinations. Alternatively, generalist interactions occur when fitness benefits do not vary across specific host-symbiont combinations. Diffuse coevolution results when the strength and direction of selection on a common partner is driven by selection from multiple mutualistic partners and/or genetic correlations across these interactions links their evolutionary trajectories. (Hougen-Eitzman and Rausher 1994; Iwao and Rausher 1997). Under diffuse coevolution, phenotypic variation across pairwise interactions will not be observed (Hougen-Eitzman and Rausher 1994; Iwao and Rausher 1997; Inouye and Stinchcombe 2011). Therefore, observing beneficial interactions that lack specificity across pairwise interactions between a range of hosts with a shared generalist symbiont indicates diffuse coevolution may underlie the interactions.

Reciprocal inoculations can be used to provide evidence of pairwise and diffuse coevolution. Between host and symbiont lineages (i.e. within species), reciprocal inoculations can be used to provide evidence of host-symbiont specificity by testing for local adaptation, where local genotypes outperform foreign genotypes in their home environments (Kawecki and Ebert 2004; Hereford 2009). Observing local adaptation demonstrates evidence of reciprocal evolutionary change between both host and symbiont, consistent with pairwise coevolution. Studies of host-parasite interactions frequently measure local adaptation to assess evidence for coevolution (Lively 1989; Ebert 1994; Thrall et al. 2002; Fischer and Foitzik 2004; Lively et al. 2004; Little et al. 2006; Greischar and Koskella 2007; Hoeksema et al. 2008; King et al. 2009), but only a few studies of mutualisms have used this approach (Hoeksema and Thompson 2007; Barrett et al. 2012; Harrison et al. 2017*a*; Rekret and Maherali 2019). Similarly, reciprocal inoculations can be used to test for evidence reciprocal evolutionary change within a genus (i.e across species) with a mutualistic partner. Observing specialization across host species with conspecific-derived symbionts demonstrates evidence of reciprocal evolutionary change between host and symbiont, consistent with pairwise coevolution, whereas observing generalist interactions may indicate a role for diffuse coevolution.

We tested the hypothesis that coevolution underlies the persistence of horizontally transmitted mutualisms. Specifically, we assessed the roles of pairwise and diffuse coevolution by testing for specialization within and across host species with their mutualistic symbionts. We tested our hypothesis by leveraging the interaction between the squash bug *Anasa tristis* and its horizontally transmitted bacterial symbiont *Caballeronia spp.* (Acevedo et al. 2021). Squash bugs that harbor *Caballeronia spp.* exhibit increased development and survival rates relative to aposymbiotic squash bugs. *Caballeronia spp.* is harbored in specialized organs, called crypts, where it grows to high titers relatively free from competition. Moreover, *A. tristis* squash bugs overlap in their geographic range with two closely related *Caballeronia*-harboring insects: *A. andresii* and *A. scorbutica*. All three insect host species and their respective *Caballeronia spp.* symbionts were isolated from their natural populations. We assessed these interactions for evidence of pairwise and diffuse coevolution using reciprocal inoculations to test for specialization within and across host species with their respective *Caballeronia* symbionts.

## Materials and Methods

### Study System

The squash bug *Anasa tristis* is an agricultural pest of cucurbit crops, such as squash and zucchini (Bonjour et al. 1990; Pair et al. 2004). Their native range includes Central America, the United States, and southern Canada (Beard 1940). Squash bug nymphs develop through five instar stages before reaching adulthood. During the second instar, squash bugs environmentally acquire the bacterial symbiont *Caballeronia* spp. (formerly called *Burkholderia* <u>spp</u>. and reclassified in 2020, but still a member of the *Burkholderiaceae* family) (Acevedo et al. 2021). These bacteria provide squash bugs with substantial fitness benefits, including increased development rate and survival to adulthood (Acevedo et al. 2021). Empirical evidence demonstrates *Caballeronia* symbionts are not passaged directly from parent to offspring through traditional vertical transmission pathways. Instead, tracking of *Caballeronia* symbionts indicates potential for symbiont transmission between co-localized individuals via the environment (Acevedo et al. 2021). The exact mechanism for symbiont transmission remains unknown but likely involves passaging through a shared environmental source, such as on or through plants. Growth within the crypt benefits *Caballeronia* spp. symbionts by allowing them to grow to high titers relatively free from competition. Most often, crypts are colonized by a single cultivable *Caballeronia* spp. strain, but coinfections are occasionally observed (Acevedo et al. 2021).

### Insect field collections and laboratory preparation

We collected *A. tristis* squash bugs and their associated *Caballeronia* symbionts across three different geographic scales. At the small geographic scale, we collected squash bugs and their associated *Caballeronia spp.* symbionts from four organic farms in Georgia, USA, separated by distances of 10 to 81 km. Field collection sites in Georgia included Woodland Gardens (WG), Front Field Farms (FFF), Crystal Organic Farm (CF), and Oxford Organic Farm (Ox). At the intermediate scale, we collected squash bugs and their associated *Caballeronia spp.* symbionts from sites across the United States, including locations in Georgia (GA), North Carolina (NC), and Indiana (IN). Squash bugs were collected from a total of four organic farms and gardens in each state. Distance between states range from 563 km to 1126 km, and distances between collection sites within each state range from 482 m to 97 km. At the large geographic scale, we collected *A. tristis* squash bugs and their associated *Caballeronia* spp. symbionts from the Eastern and Western United States. Western bugs were collected from sites in Arizona, USA, and Eastern bugs were collected from sites in Georgia, USA (2789 km between states). For specialization assays across host species, we collected *A. tristis*, *A. andresii*, and *A. scorubtica* hosts from four sites in Gainesville, Florida separated by a distance of 1.6 to 14.5 km. Following field collections, bugs were returned to the lab, scanned for ectopic parasites, and allocated for use in either the establishment of experimental populations or symbiont isolation. To establish populations, bugs were placed on yellow crookneck squash plants in environmental chambers at 21 °C, 60% humidity, and a 16/8 h day/night cycle. Bugs from each collection site were placed separately on two to three plants, each housing two mating pairs each for a total of thirty to fifty bugs per state.

For specialization assays across host species, we collected *A. tristis*, *A. andresii*, and *A. scorubtica* hosts from four sites in Gainesville, Florida separated by a distance of 1.6 to 14.5 km. Insects were returned to the lab, scanned for ectopic parasites, and allocated for use in either establishment of experimental populations or symbiont isolation. Insects used to produce progeny for inoculations were placed on squash plants in environmental chambers under the same conditions described previously. Insects from each species were placed on five plants with conspecifics, each housing three to four mating pairs, for a total of thirty to forty bugs per species.

### Genetic analyses of *Caballeronia* symbionts

Testing for local adaptation and genetic specificity requires there to be phenotypic and genetic variation across the hosts and symbionts involved in the experiments (Gandon and Van Zandt 1998). For the *Caballeronia* symbionts, we assessed whether there was genetic variation in the isolates used for experiments and additional isolates from the same populations in three ways. We first used traditional sanger sequencing of 16s rRNA to identify *Caballeronia* symbionts isolated from field collected squash bugs. We then used whole genome sequencing to gain greater insight into the genetic variation present in the symbiont populations. Finally, we used high throughput 16s rRNA sequencing of whole crypt communities to gain greater insight into the distribution of *Caballeronia* variants across individuals and populations.

To isolate individual symbionts for local adaptation assays, we dissected the crypts of one to ten squash bugs per each collection site, depending on the availability of field collected bugs from each site, and isolated bacterial symbionts (n = 34 GA bugs + 16 NC bugs + 26 IN bugs = total 76 bug dissections). We isolated individual symbionts from each host species by dissecting the crypts from bugs that were collected in Gainesville, Florida (n = 44 *A. tristis* bugs + 28 *A. andresii* + 16 *A. scorbutica*). For each crypt, half was placed in 1x PBS and homogenized by crushing. The other half was stored in ethanol (99%) for later culture-independent assessment of the bacterial community composition. Crypts placed in 1x phosphate buffered saline (PBS) were crushed, dilution plated onto LB agar, and grown at 28 °C for 48-72 hours. Resulting colonies that morphologically resembled *Caballeronia spp.* were individually selected from each plate and stored in glycerol (15%) at −80 °C. In general, bacteria morphologically similar to *Caballeronia* were the predominant bacteria on the plates.

For sanger sequencing, bacteria stored in glycerol were revived by streaking onto LB agar and grown at 28 °C for 48 hours. DNA from each bacterium was extracted by boiling at 95 °C (Dashti et al. 2009). Extracted DNA was amplified by polymerase chain reaction (PCR) using general 16s ribosomal DNA bacterial primers, including the forward primer 27F (5’- AGAGTTTGATCMTGGCTCAG-3’) and reverse primer 1492R (5’- GGTTACCTTGTTACGACTT-3’) (Lane 1991). PCR amplifications were performed with an initial 4 minutes of denaturing at 94 °C, followed by 36 cycles of denaturing for 30 seconds at 94 °C, annealing for 30 seconds at 55 °C, extending for 1 minute at 72 °C, and a final 1-minute extension at 72 °C. PCR products were purified using a Qiagen QIAquick PCR purification kit and protocol. Samples were sanger sequenced, and the resulting sequences were assembled in DNASTAR SeqMan Pro. Aligned sequences were run through the NCBI NIH BLAST nucleotide database for identification. We identified a total of 26 *Caballeronia* strains from bugs isolated in Georgia, North Carolina, and Indiana (GA = 14; NC =4; IN = 6). We identified a total of fifteen strains from bugs of each host species isolated in Florida (*A. tristis* = 6, *A. scorbutica* = 4, *A. andresii* = 5). Failure to isolate *Caballeronia* strains likely resulted from the limitations of culture-based methods, rather than the absence of *Caballeronia* within squash bug crypts, as shown by 16s rRNA sequencing of the whole crypt community (see below). Sequences identified as *Caballeronia* spp. were aligned using DNASTAR MegAlign Pro, and we calculated the percent identity of the 16s gene across strains at each scale. We selected the most genetically dissimilar strains at each geographic scale and from each host species for reciprocal inoculations.

We conducted whole genome sequencing for *Caballeronia* strains isolated from *A. tristis* in Georgia (n = 13 strains), Indiana (n = 6 strains), and North Carolina (n = 4 strains). We also conducted whole genome sequencing from strains isolated from *A. tristis* (n = 6 strains), *A. andresii* (n = 4 strains), and *A. scorbutica* (n = 5) hosts collected in Florida. Bacterial strains were revived from glycerol by streaking onto LB agar and grown for 48 hours at 28 °C. Individual colonies were selected and grown in LB overnight with shaking at 28 °C. DNA was extracted from bacteria in liquid cultures using a Qiagen DNeasy blood and tissue kit. DNA quality was assessed using gel electrophoresis and quantity was measured using a Nanodrop spectrometer. Genomic sequences were obtained using paired-end Illumina Mi-seq whole genome sequencing. Reads were assembled *de novo* using the SPAdes genome assembler, version 3.15.3 (Bankevich et al. 2012). Contigs were ordered against a reference genome using Mauve contig mover (Rissman et al. 2009). Genome assembly completeness was assessed using BUSCO (Simão et al. 2015). Multiple genome alignment and comparative genome analysis were conducted using anvi’o (Eren et al. 2021). We performed pangenome analysis to compare amino acid sequences and gene cluster presence and absence across symbiont strains using the program DIAMOND (Buchfink et al. 2014). Evolutionary relationships were estimated using single-copy core genes clusters for phylgenomic analysis. We assessed genetic variation across genomes by computing the average nucleotide identity across genomes using PyANI (Pritchard et al. 2016).

For 16s rRNA-based analyses of entire microbial communities within individuals’ crypts, the crypts previously stored in ethanol at −80 °C were thawed and rinsed in 1x PBS. DNA was extracted using the Lucien MasterPure Complete DNA and RNA Purification kit protocol and reagents. Crypts stored in ethanol following insect dissection were rinsed in 1x PBS. DNA was purified using the ZYMO OneStep PCR Inhibitor kit and subsequentially quantified using a Thermo Fisher Scientific NanoDrop One UV Spectrophotometer. Samples for high-throughput sequencing were selected based on successful amplification of the V3-V4 region of 16S rRNA gene via PCR. We used 341f/785r primers, as described in Klindworth et al. 2013, for DNA amplification. PCR reagents, protocol, and thermocycling conditions adhered to the New England BioLabs *Taq* PCR kit. PCRs were performed on Eppendorf Mastercycler. Gel visualizations were run on Agilent’s 4200 TapeStation using the D1000 Screentape. Samples with clear bands were selected for Illumina Miseq sequencing. Statistical comparisons of crypt communities across host populations were performed using PERMANOVA.

### Genetic analyses of *Anasa tristis*

The population structure of *A. tristis* was assessed using restriction site-associated DNA sequencing (RAD-seq). We extracted DNA from 95 bugs collected from Georgia (n=50), Indiana (n=20), and North Carolina (n=20). DNA was extracted using either Omega Bio-tek E.Z.N.A Insect DNA kits or Qiagen DNeasy blood and tissue extraction kits. DNA quality control, library prep and sequencing were performed by CD Genomics. DNA quality and purity was assessed using gel electrophoresis and nanodrop spectrometry. DNA was quantified using a Qubit 2.0 flurometer. Library construction was performed using an Illumina TruSeq Nano DNA Sample Prep Kit. High throughput sequencing was performed using the Illumina novaseq6000 platform with a read length of 150 bp at each end. Several samples across host populations produced low quality reads that were excluded from analysis. Final bioinformatic analysis included comparison of 44 GA squash bugs and 19 squash bugs from both IN and NC. Reads were mapped to a reference genome using BWA. SNP and InDel detection and analysis were performed using GATK. We built a phylogenetic tree using maximum likelihood to estimate evolutionary relationships. Variation across populations was assessed using principal component analysis, and genetic divergence between populations was assessed by measuring pairwise Fst (Holsinger and Weir 2009).

### Experimentally assessing host-symbiont specificity by testing for local adaptation

We established squash bug lineages for local adaptation assays, as described above. Field collected squash bugs reproduced in the environmental chambers, and we performed experiments using the second (F2), third (F3) and fourth (F4) generation progeny to reduce environmental and maternal effects. Reciprocal inoculations at the smallest geographic scale included progeny from the F2 generation, the intermediate scale included a mixture of progeny from the F2 and F3 generations, and the largest scale included a mixture of progeny from the F3 and F4 generations for Eastern *A. tristis* and the F2 progeny for Western *A. tristis*.

We selected a single symbiont strain from each collection site for reciprocal inoculations at the smallest geographic scale (GAWG2-4, GACF4, GAFFF3, and GAOX1). We selected a single strain from each state for reciprocal inoculations at the intermediate scale (GACF4, INML1, and NCF4) and a single strain for each state at the large geographic scale (GACF4 and AZ1). Strains for the small and intermediate scale were selected as described above, such that reciprocal inoculations were conducted using the most genetically dissimilar strains at each geographic scale.

Prior to inoculations, frozen *Caballeronia spp.* samples were revived as described above. Liquid feeding solutions for symbiont inoculations were prepared as described in Acevedo et al. 2021. Specifically, liquid cultures were prepared and incubated overnight at 28 °C with shaking. Overnight cultures were diluted 1:5 in LB and incubated at 28 °C with shaking for two hours. Bacterial feeding solutions (10mL) were prepared by diluting the two-hour liquid cultures with sterile molecular water to ∼2×10^7^ cells/mL. Blue dye (1%) was added to each solution to allow for visual confirmation of the feeding solution in squash bug guts. Feeding solutions were poured into 35mm Petri dishes, and a cotton dental swab was placed in each dish for squash bugs to feed from. Plates were wrapped in parafilm to prevent spilling or squash bug drowning.

For reciprocal inoculations at the small geographic scale, squash bug eggs from each mating pair for a given collection site were pooled. Eggs were surface sterilized with ethanol (70%) and bleach (10%) and returned to environmental chambers. Emerging first instar nymphs were fed surface sterilized zucchini or squash. At second instar, nymphs were starved for seven to nine hours. Symbiont inoculations were performed such that squash bugs from each collection site were provided with bacterial feeding solutions from each collection site (*i.e*. one sympatric and three allopatric combinations per host population). After 24 hours, feeding solutions were removed. Thirty nymphs from each treatment were divided into plastic vented containers housing five bugs each and fed zucchini or squash. Due to limitations in egg production, inoculations using each symbiont strain occurred separately over four consecutive weeks. This procedure was repeated until all symbionts were fed to hosts from each population. In total, hosts from each collection site were inoculated with bacteria from each collection site for a total of four sympatric (n = 30 bugs/treatment x 4 sympatric treatments = total of 120 bugs receiving sympatric bacteria) and twelve allopatric inoculations (n = 30 bugs/treatment x 12 allopatric treatments = total 360 bugs receiving allopatric bacteria). Aposymbiotic controls were established by feeding nymphs water according to the protocol above. Twenty-five hosts from each population were provided water solutions (n = 25 hosts/populations x 4 populations = total of 100 water control bugs). A small number of bugs died across host populations following water inoculations, and fitness measurements were conducted for 20-25 hosts per population (WG = 20 bugs; FFF = 25 bugs; Ox = 24 bugs; CF = 21 bugs; n = 90 total water control bugs). Host survival and development rate were assessed every other day until bugs reached adulthood.

Inoculations with each bacterial symbiont were performed asynchronously, which could conflate symbiont effects with replicate effects. To address this limitation, we later tested explicitly for symbiont effects by inoculating nymphs from a single host population with bacteria from all four collection sites on the same day. Tests for symbiont effects were performed for the CF and FFF host populations, which were the only host populations from which enough eggs could be collected. Eggs were sterilized and squash bugs were inoculated according to the procedure described previously. Bacteria from each collection site were fed to 15 to 20 squash bugs from each of the two populations. Host fitness was measured as described above.

Symbiont fitness was measured as the number of colony forming units (CFUs) per adult squash bug crypt. Adult squash bugs were anesthetized using carbon dioxide and surface sterilized in ethanol (70%). Crypts were manually dissected, placed in 1x PBS, crushed, and serially diluted. Spread plates were prepared on LB agar and grown at 28 °C for 48 hours. The number of colonies per plate was counted. Observation of multiple colony morphologies was noted, and only colonies with morphological characteristics of *Caballeronia spp.* were counted. Bacteria morphologically typical of *Caballeronia* were the predominant bacteria on the plates.

For reciprocal inoculations at the intermediate geographic scale, eggs from all collection sites for a given state were pooled. Egg sterilization and bacterial inoculations were performed as described above. For each inoculation, second instar nymphs from each state (IN, GA, and NC) were provided bacteria isolated from each state (*i.e*. one sympatric and two allopatric combinations per inoculation for the intermediate geographic range). Twenty-five nymphs from each treatment were placed in groups of five in plastic vented containers and fed zucchini or squash (total of five containers per treatment). This procedure was repeated three times for full reciprocity for a total of three sympatric combinations (n = 25 bugs/treatment x 3 sympatric treatments = total of 75 bugs receiving sympatric bacteria) and six allopatric combinations (n = 25 bugs/treatment x 6 allopatric treatments = total of 150 bugs receiving allopatric bacteria). Aposymbiotic water controls were produced for each host population as described above (n = 15 bugs/populations x 3 populations = total of 45 water control bugs). Following the establishment of these reciprocal inoculations, the inoculations were replicated a second time so that all host populations were fed bacteria from all collection sites on the same day, allowing distinction between symbiont versus replicate effects. In this case, twenty bugs from each collection site were inoculated with bacteria from each collection site for a total of three sympatric (n = 20 bugs/treatment x 3 sympatric treatments = total 60 bugs receiving sympatric bacteria) and six allopatric inoculations (n = 20 bugs/treatment x 6 allopatric treatments = total of 120 bugs receiving allopatric bacteria). Across the two replicates, a total of 135 bugs received sympatric bacteria, and 270 bugs received allopatric bacteria. Due to the global COVID-19 pandemic and lab closures, squash bugs from each replicate were removed from environmental chambers and kept at ambient temperature (27 °C), on days 40 and 19 of each replicate experiment, respectively, in order to keep the experiment going. Host fitness was measured over the course of the experiment as described previously. Due to the pandemic and lab closures, we were unable to assess symbiont fitness for reciprocal inoculations at the intermediate geographic scale.

Reciprocal inoculations at the large geographic scale were performed using *A. tristis* collected from Arizona and Georgia. Egg collection, sterilization, and feeding solutions were prepared as described previously. Bugs from each state were provided bacteria from each state for a total of two sympatric (n = 30 bugs/treatment x 2 sympatric treatments = total 60 total bugs receiving sympatric bacteria) and two allopatric treatments (n = 30 bugs/treatment x 2 sympatric treatments = total 60 total bugs receiving allopatric bacteria). Inoculated bugs were placed in groups of five in plastic vented containers and fed zucchini or yellow crookneck squash. Host survival and development rate were recorded as described previously. Symbiont fitness was measured at this geographic scale as described for the small geographic scale.

### Testing for specialization between host species and *Caballeronia spp*. strains

The native geographic range of *A. tristis* overlaps with two closely *Caballeornia*-harboring sister species, *A. andresii* and *A. scorbutica* (Acevedo et al. 2021). In the field, these species are observed inhabiting the same plants, indicating the potential for symbiont sharing between species. We tested for specialization between these overlapping species with their respective *Caballeronia spp.* strains by performing reciprocal inoculations.

Field collected squash bugs were used to establish laboratory colonies and bacterial strains for reciprocal inoculations, as described previously. We randomly selected a single bacterial strain isolated from each host species for reciprocal inoculations: AAF181, ATF2731, and ASM285. Reciprocal inoculations were performed so that each host species was inoculated with bacteria isolated from each host species for all possible combinations of host and bacteria (*i.e.* one conspecific-derived symbiont and two heterospecific-derived symbionts for each host species x 3 host species = total of three conspecific-derived symbiont inoculations and six heterospecific-derived inoculations). For *A. tristis*, we inoculated a total of 25 bugs per treatment (n = 25 bugs receiving conspecific-derived symbionts + 50 bugs receiving heterospecific-derived symbionts = 75 total bugs). For *A. andresii*, we inoculated a total of 25 bugs per treatment (n = 25 bugs receiving conspecific-derived symbionts + 50 bugs receiving heterospecific-derived symbionts = total 75 bugs). For *A. scrobutica*, we inoculated a total of 20 bugs per treatment (n = 20 bugs receiving conspecific-derived symbionts + 40 bugs receiving heterospecific-derived symbionts = 60 total bugs). We prepared bacterial feeding solutions and measured host and symbiont fitness as described for the local adaptation assays.

### Statistical analysis

For reciprocal inoculations within species, we performed used cox proportional hazard models to test whether host survival varied based on the following main effects included in the model: host origin, symbiont origin, or in response to an interaction between host and symbiont (Table 1). Death was considered an event, and bugs were censored once reaching adult. Occasionally bugs were accidentally killed while collecting data, and these bugs were censored. For the intermediate scale, we repeated reciprocal inoculations two times, and the effect of inoculation block was included as a fixed effect for both survival and development rate analyses. We also performed cox proportional hazard models using the survival package in R to test whether host development rate to adult varied in response to the following main effects: host origin, symbiont origin, or an interaction between host and symbiont origin (Table 1). We chose to perform analysis for the rate to adult because variation across treatments was greatest to this stage, and bugs reach reproductive maturity at adult, making the rate to adult the best measure of host fitness. We considered reaching adulthood as an event, and bugs that died before reaching adulthood were censored. If we detected an interaction between host and symbiont origin for survival or development rate to adult, we performed contrasts using the emmeans package in R to test for an effect of sympatric versus allopatric combinations of host and symbiont.

**Table 1.**
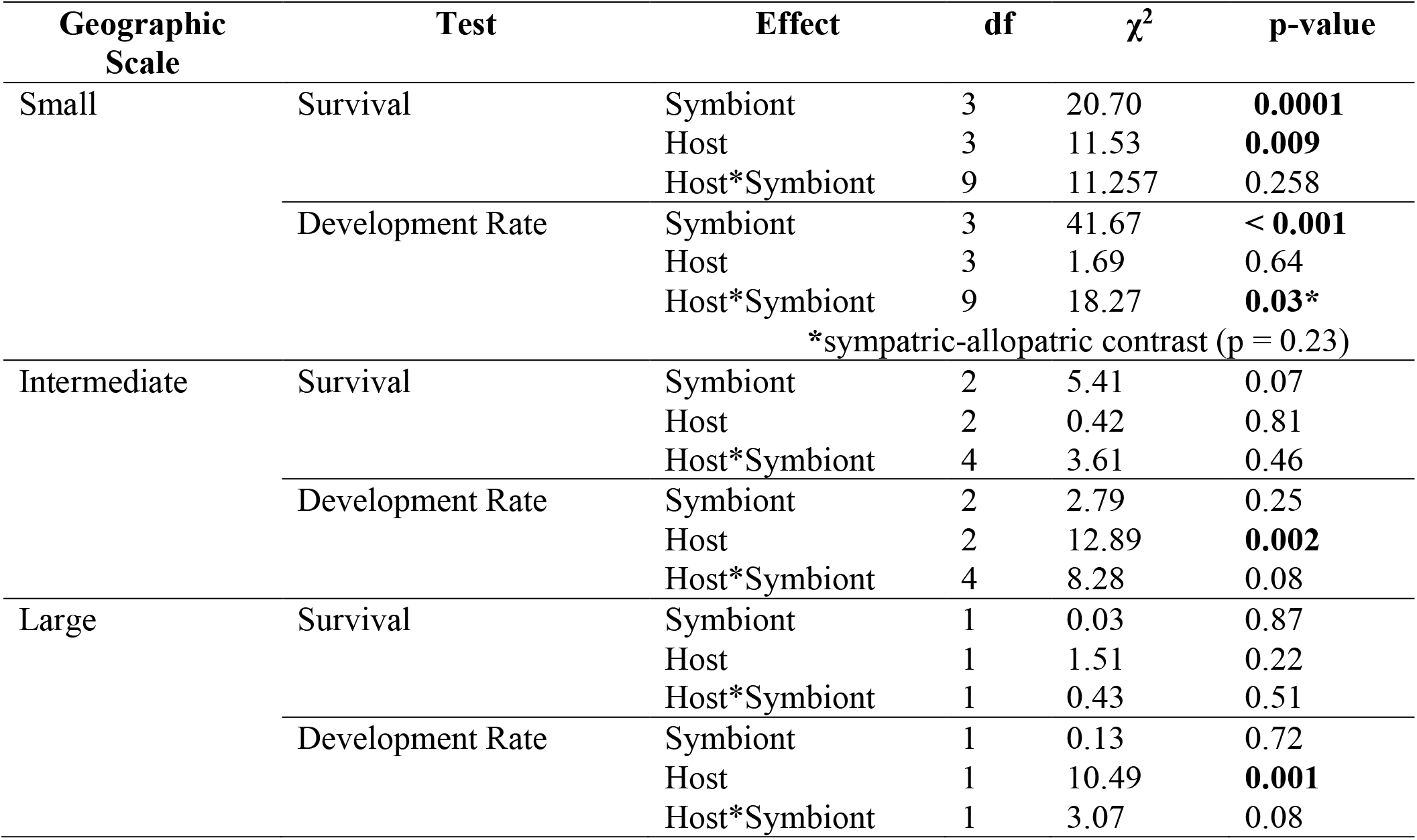
Statistics for host survival and development rate for local adaptation reciprocal inoculations. We performed cox proportional hazard models to test for effects of geographic origin of host and symbiont and an interaction between host and symbiont geographic origin on host time to death and time to adult. If a significant interaction was observed, we performed a linear contrast to test whether host fitness varied in response to sympatric versus allopatric symbionts.

For reciprocal inoculations across species, we used a cox proportional hazard analysis to test whether host survival varied in response to symbiont origin, which was included as a main effect (Table 2). We considered death an event, and bugs that reached adult were censored. We used a cox proportional hazard model to test whether host rate of development to adult varied in response to symbiont origin, which was included as a main effect (Table 2). Analyses for each host species were performed separately because difference in the rate of survival and development are likely intrinsic to each separate species. We considered reaching adulthood as an event, and bugs that died before reaching adulthood were censored.

**Table 2.**
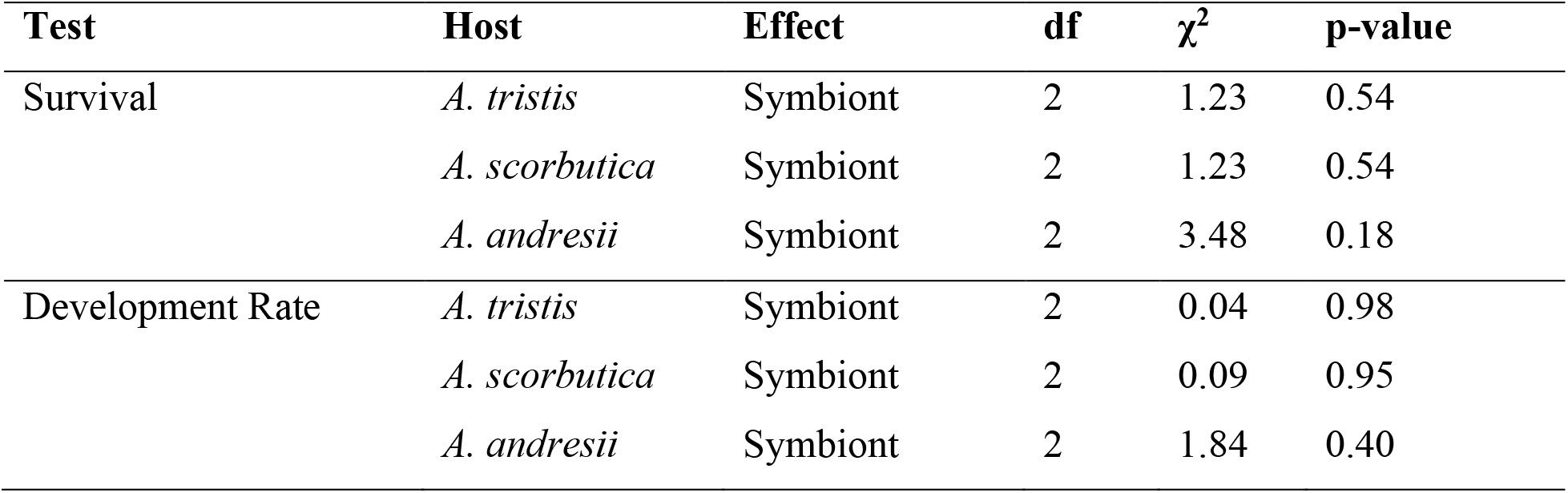
Statistics for host development rate to adult and survival for reciprocal inoculations between three host species and symbionts derived from conspecific versus heterospecific hosts. We performed cox proportional hazard models to assess rate of development to adult and survival.

| Test | Host | Effect | df | $\chi^2$ | p-value |
| --- | --- | --- | --- | --- | --- |
| Survival | *A. tristis* | Symbiont | 2 | 1.23 | 0.54 |
|  | *A. scorbutica* | Symbiont | 2 | 1.23 | 0.54 |
|  | *A. andresii* | Symbiont | 2 | 3.48 | 0.18 |
| Development Rate | *A. tristis* | Symbiont | 2 | 0.04 | 0.98 |
|  | *A. scorbutica* | Symbiont | 2 | 0.09 | 0.95 |
|  | *A. andresii* | Symbiont | 2 | 1.84 | 0.40 |

For analysis both within and across species, we further assessed host survival using a Chi-squared test to assess the proportion of hosts surviving to adulthood in sympatric versus allopatric combinations of host and symbiont. We used a quasipoisson distributed generalized linear model to test whether symbiont fitness (logCFU/crypt) varied in response to the following main effects: host origin, symbiont origin, or an interaction between host and symbiont origins. Water was excluded from models comparing the effect of host and symbiont origin on partner fitness, and we compared the effect of receiving a symbiont versus water treatment on host development rate and survival using the same statistical tests mentioned above. Models were fit using R, version 4.1.0.

## Results

### Geographic genetic variation is observed across *A. tristis* host and symbiont populations

We performed genetic analysis to test for spatial structure and underlying genetic variation across *A. tristis* host and symbiont populations at the intermediate geographic scale. In general, host and symbiont populations vary genetically across their geographic range, suggesting the potential for local adaptation. We observed moderate differentiation between the IN population with both the host populations from GA (Fst = 0.070) and NC (Fst = 0.075). These results demonstrate moderate spatial structure and genetic variation, such that *A. tristis* hosts from GA have diverged from IN and NC populations.

We observed similar geographic variation across symbiont strains. Based on analyses of whole genome sequence data, we observed little within-population variation for symbionts isolated from IN and NC (ANI ∼0.99; Figure 2). In contrast, we observed higher genetic variation within the GA symbiont population (ANI = 0.82-0.99; Figure 2). In general, symbionts isolated from NC and IN shared high average nucleotide identity with each other (ANI = 0.90 – 0.99; Figure 2, S1). On average, strains from GA shared higher average nucleotide identity with strains from NC than IN, but average nucleotide identity across all pairwise combinations of NC and GA strains varied broadly (ANI = 0.82 – 0.99; Figures 3, S1). This variation was driven by several GA strains that were genetically diverged from all other strains, forming their own monophyletic clade (Figure 2).

**Figure 1.**
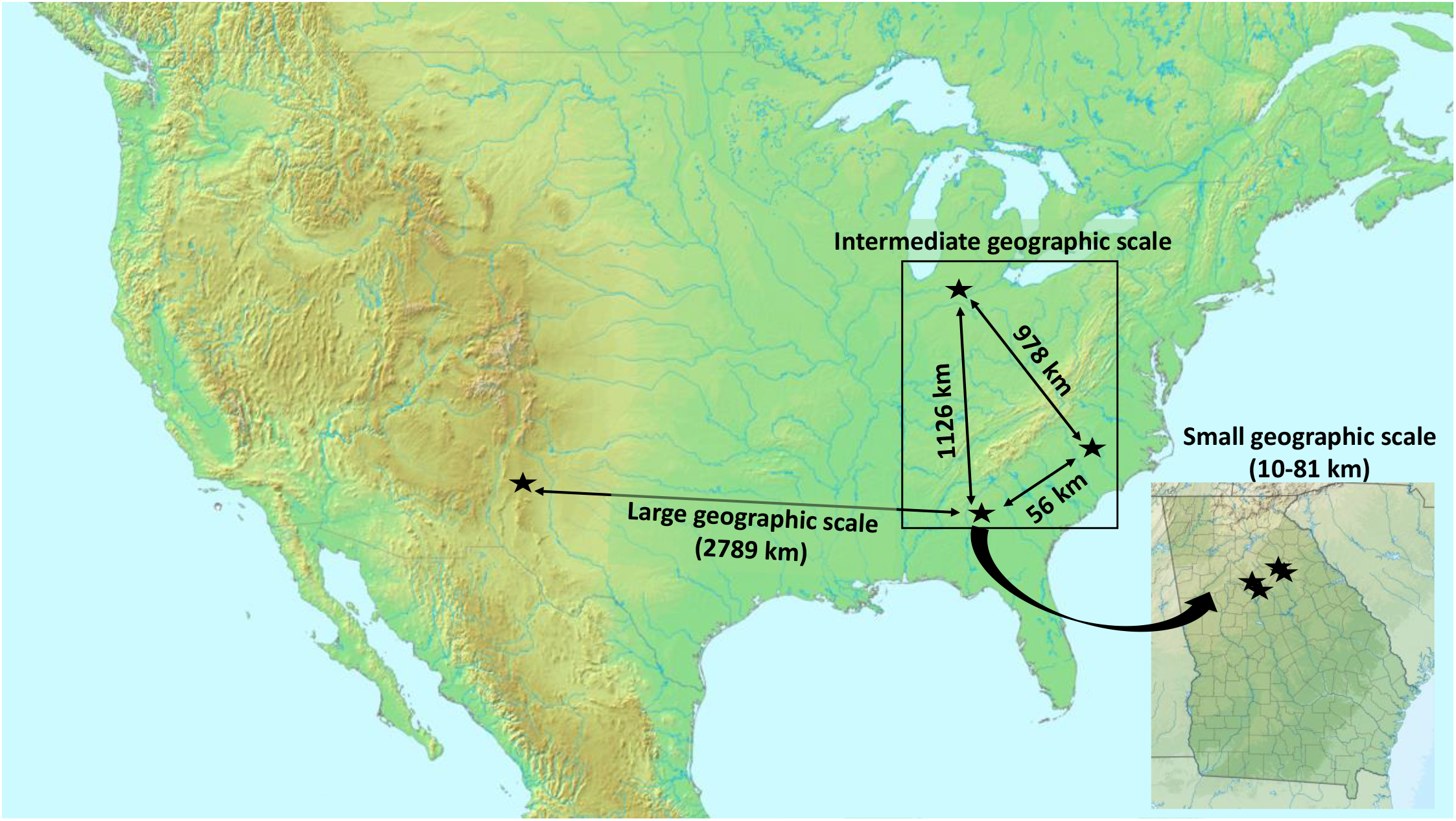
Geographic scales for local adaptation assays. *A. tristis* hosts were collected from sites across three geographic scales: small, medium, and large. At the small geographic scale, bugs were collected from four sites in Georgia, USA. At the intermediate scale, we collected *A. tristis* from three different states: Georgia, Indiana, and North Carolina. At the large spatial scale, Eastern USA *A. tristis* were collected in Georiga sites and Western USA *A. tristis* were collected in Arizona.

**Figure 2.**
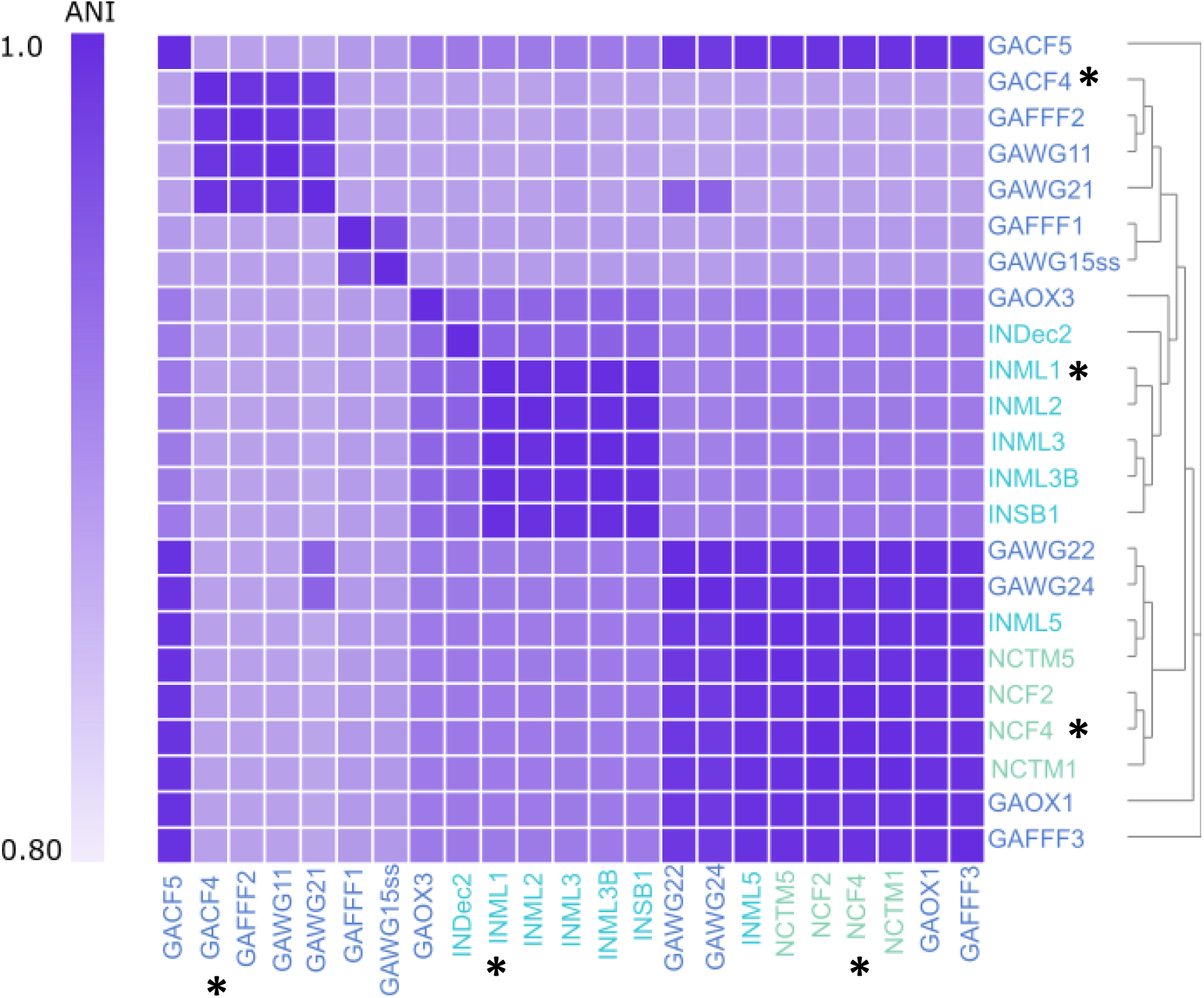
Heat map showing the genetic distance across symbiont strains isolated from *A. tristis* from Georgia (GA; purple), Indiana (IN; blue), and North Carolina (NC; green). Genetic distance was calculated using average nucleotide identity (ANI), which is a measure of the nucleotide similarity across coding regions. ANI varied from 0.82 to 1.0, with the largest difference resulting when comparing IN and GA symbionts. Far right, evolutionary relationships between strains were assessed using phylogenomics. Strains GACF4, NCF4, and INML1, asterisks, were used for experimental reciprocal inoculations.

**Figure 3.**
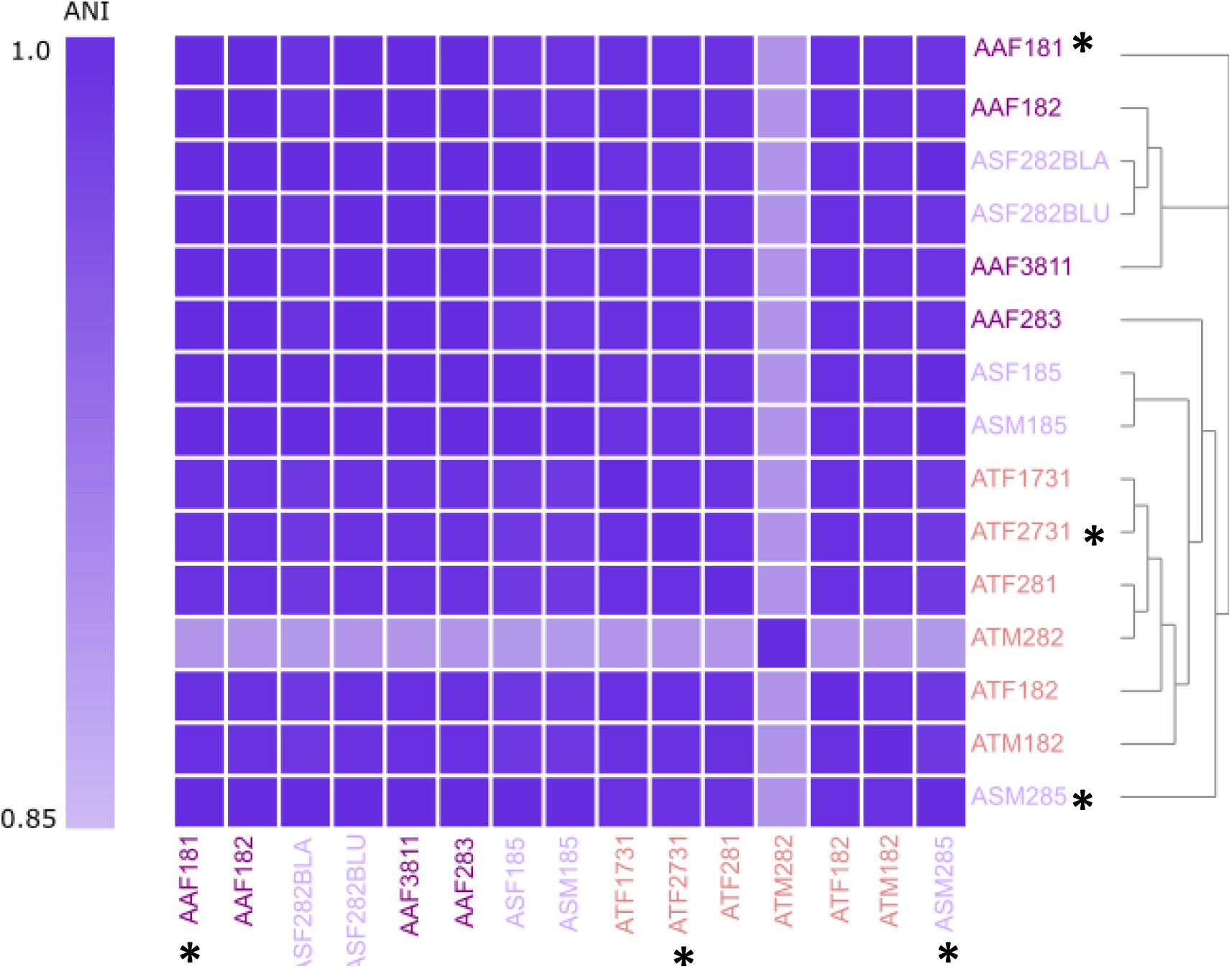
Heat map showing the genetic distance across symbionts strains isolated from *A. tristis* (AT; pink), *A. andresii* (AA; dark purple), and *A. scorbutica* (AS; light purple). All hosts and their associated symbionts were collected in Florida, USA. Genetic distance was calculated using average nucleotide identity (ANI), which varied from 0.85 to 0.99. We observed little variation across strains isolated from heterospecific hosts, with only one strain (ATM282) exhibiting substantial variation from other strains. Far right, evolutionary relationships between strains were assessed using phylogenomic analysis. Strains ASM285, AAF181, and ATF2731, asterisks, were used for reciprocal inoculations to test for specialization between host species and symbiont strain.

We further assessed geographic symbiont strain variation using 16s rDNA-based community analysis, which allows for comparison of both cultivable and uncultivable symbiont strains within the host crypts. We observed a similar effect of geographic origin on crypt *Burkholderiaceae* community composition (richness and evenness) (F_2,36_ = 3.17, p = 0.003; Figures 4, S3) as we did for whole genome-based analyses. Specifically, the *Burkholderiaceae* community composition of crypts isolated from NC and IN bugs were more similar than those isolated from GA bugs (F_1,37_ = 2.41, p = 0.033; Figures 4, S3). This is the same pattern observed across whole genome analysis of the cultivable symbiont strains across GA, IN, and NC.

**Figure 4.**
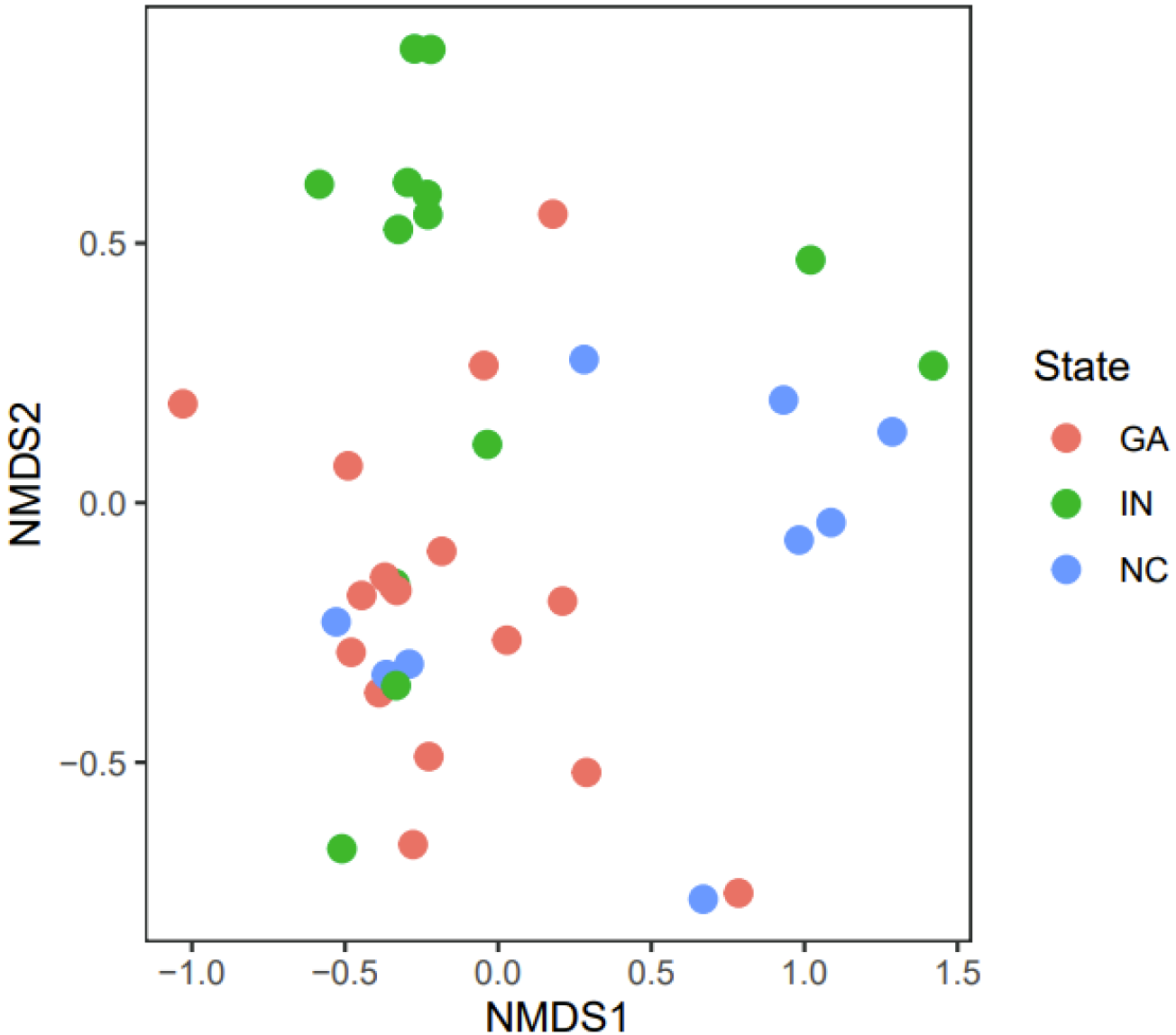
NMDS plot demonstrating variation within the *Burkholderiaceae* crypt communities of squash bugs isolated across the intermediate geographic range: Georgia (GA; pink), Indiana (IN; green), and North Carolina (NC; blue). Each dot represents all OTUs within the *Burkholderiaceae* (bacterial family including *Caballeronia spp.*) for a single crypt isolated from a bug. Crypt community composition varied across the intermediate geographic range (F_2, 36_ = 3.17, p = 0.002). Differences between strains are driven by differences in GA crypt community composition relative to those of IN and NC (F_1,37_ = 2.47, p = 0.037).

### Genetic variation in symbiont strain is not observed across host species

Using whole genome sequencing, we observed little nucleotide variation across symbiont strains isolated from *A. tristis*, *A. scorbutica*, and *A. andresii* hosts. All strains in this analysis were isolated from *A. tristis*, *A. scorbutica*, and *A. andresii* hosts collected in Florida, USA, eliminating the potential for geographic based genetic variation. All strains shared high average nucleotide identity (ANI = 0.99; Figure 3, S2), except for one strain isolated from an *A. tristis* host that differed from all other strains (ATM282; ANI = 0.85; Figure 3, S2). Symbiont strains isolated from heterospecific hosts do not vary from one another in their average nucleotide identity, suggesting specialization between strain and host species as an unlikely outcome.

### Local adaptation is not observed between *A.tristis* hosts and their symbionts

We tested for specificity consistent with pairwise coevolution by measuring local adaptation between hosts and symbionts across a small, intermediate, and large geographic scale. We quantified host response to their sympatric symbiont and a range of allopatric symbionts by measuring host survival and development rate to each life stage. If pairwise coevolution contributes to the maintenance of this interaction, we predicted that we would observe specificity between hosts and their local *Caballeronia spp*. symbiont strains. Specifically, we predicted we would observe faster development rates and higher survival for squash bugs, and higher growth within crypts for symbionts, paired with their sympatric versus allopatric partners. For each reciprocal inoculation, we designated bugs to an aposymbiont water control treatment. Across all analyses both squash bugs receiving water exhibited slower development rate and reduced survival compared to those that received a symbiont, consistent with Acevedo et al. 2021. These individuals were not included in subsequent analyses.

#### Tests for local adaptation at the small geographic scale

At the small geographic scale, we detected a significant effect of the geographic origin of host (χ^2^ = 11.55, df = 3, p = 0.009; Table 1) and symbiont (χ^2^ = 20.70, df = 3, p = 0.0001; Table 1) on host survival. We did not detect a significant interaction between geographic origin of host and symbiont for host survival (Table 1; Figures 5, S4). Moreover, there was no significant difference in the proportion of hosts that survived to adult for sympatric and allopatric combinations of host and symbiont (Table 1; Figures 5). We observed a significant effect of the geographic origin of symbiont (χ^2^ = 41.67, df = 3, p < 0.001) and a significant interaction between host and symbiont geographic origin (χ^2^ = 18.27, df = 9, p = 0.03) for host development rate to adult (Table 1; Figures 5, S4). Because we observed a significant interaction between host and symbiont, we hypothesized that differences in development rate to adult result from local adaptation. To test this hypothesis, we contrasted the rate of development for hosts paired with sympatric versus allopatric symbionts. Host development rate did not vary in response to sympatric versus allopatric symbionts, so we did not find support for our hypothesis of local adaptation. For symbiont fitness, we measured the number of CFUs per crypt of surviving adult squash bugs (Figures 5, S5). We detected a significant effect of host (F_3,169_ = 3.0913, p = 0.02878) and symbiont (F_3,166_ = 20.2003, p < 0.001) (Figure S5). However, we did not observe a significant interaction between geographic origin of host and symbiont for CFUs per squash bug crypt (F_9,157_ = 10.04, p > 0.11) (Figures 5, S5). Taken together, these results indicate that despite phenotypic variation in the effect of the geographic origin of host and symbiont for host fitness, local adaptation has not evolved at this geographic scale.

**Figure 5.**
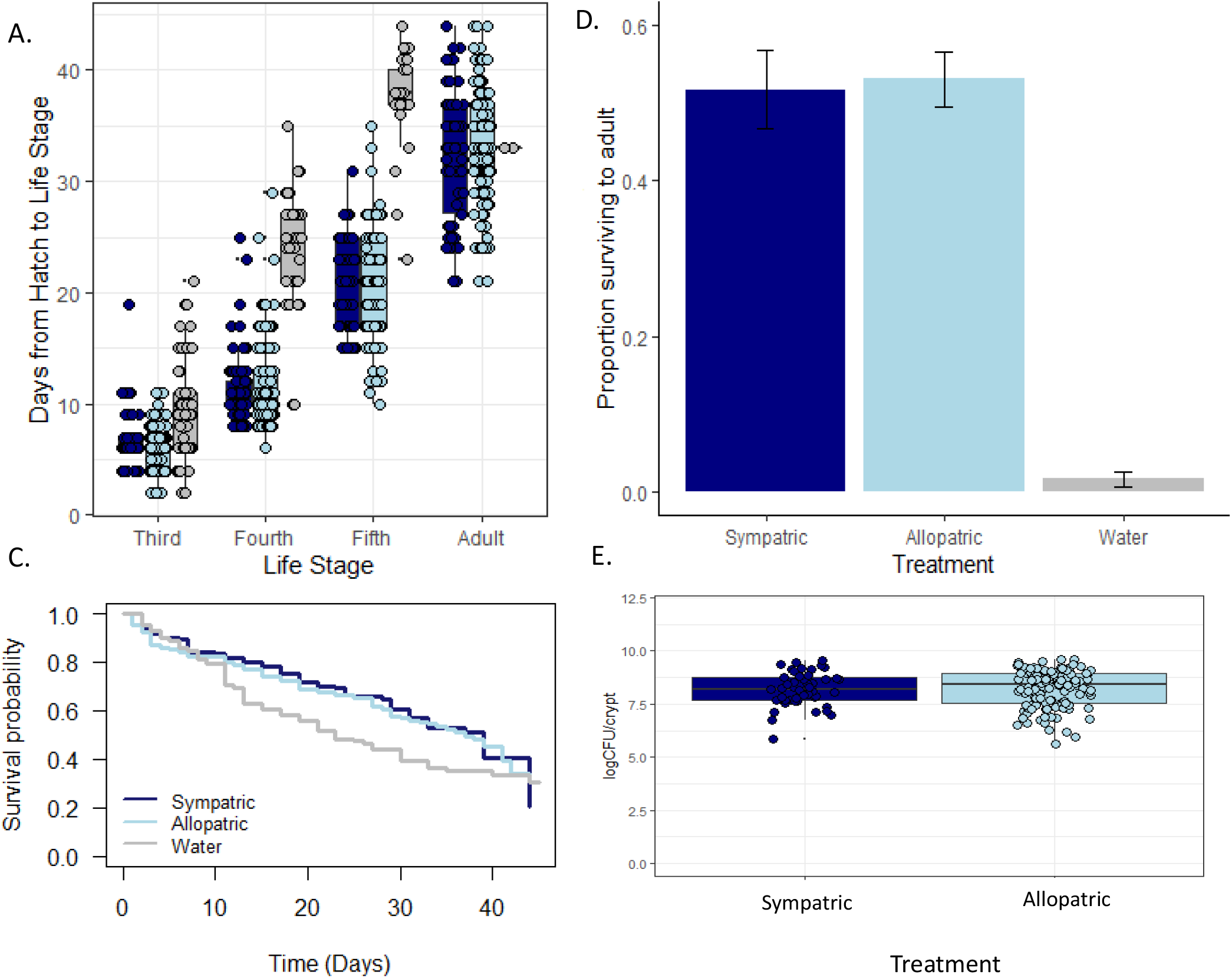
Host and symbiont fitness for reciprocal inoculations to test for local adaptation at the small geographic scale. Figure 5A shows host development rate across all developmental life stages. We performed a cox proportion hazard model to determine whether host rate of development to adult varied in response to host origin, symbiont origin, or an interaction between host and symbiont origin. We observed a significant effect of symbiont (p < 0.001) and an interaction between host and symbiont (p = 0.03), which was not driven by differences in the effect of sympatric versus allopatric symbionts on host fitness (p = 0.23). Hosts receiving water developed slower than those that received a symbiont across all life stages. See figure S4 for host development rate across all pairwise combinations of host and symbiont. Figure 5B shows the proportion of bugs surviving to adult across experimental treatments. The proportion of bugs surviving to adult did not vary across sympatric and allopatric treatments (p = 0.75) but was significantly lower for bugs receiving water versus those receiving a symbiont (p < 0.001). Figure 5C shows the survival curves across symbiont treatments (see figure S4 for survival curves across all pairwise combinations of host and symbiont). We performed a cox proportional hazard model to determine whether host survival varied in response to host origin, symbiont origin, or an interaction between host and symbiont origin. We observed and effect of symbiont (p = 0.0001) and host (p = 0.009) for host survival. Figure 5D shows the effect of treatment on symbiont fitness (logCFU/crypt) (see figure S6 for each pairwise combination of host and symbiont). We observed no effect of host on symbiont fitness.

Because reciprocal inoculations were not performed synchronously, we predicted the observed effects of host and symbiont may reflect variation across replicate rather than variation across strains. To further assess whether geographic origin of host and symbiont affects host survival and development rate, we repeated reciprocal inoculations for two host populations (CF and FFF), such that each symbiont strain was synchronously provided to each host population. When inoculations were performed synchronously, we did not detect an effect of geographic origin of host nor symbiont for host survival (Table S1; Figure S6). However, we did observe an interaction between host and symbiont geographic origin for host survival (χ^2^ = 8.92, df = 3, p = 0.03). We contrasted the survival of hosts paired with sympatric versus allopatric symbionts to test whether this interaction resulted from local adaptation. We observed no difference in host survival when paired with a sympatric versus an allopatric symbiont (p = 0.90), providing no support for local adaptation between host and symbiont. When inoculations were performed synchronously, we observed no effect of symbiont geographic origin, host geographic origin, nor an interaction between host and symbiont geographic origin, for rate of development to adult (Table S1; Figure S6). These results indicate that previous variation in the effect of symbiont for host fitness likely resulted from variation across replicate rather than as a result of variation across sites. Moreover, these results are consistent with those obtained previously and provide further support for a lack of local adaptation between host and symbiont at this small geographic scale.

#### Tests for local adaptation at the intermediate geographic scale

Gene flow can dilute the strength of local selection by decreasing opportunities for conserved interactions and reciprocal selection between host and symbiont lineages across generations. Therefore, we tested for genetic specificity at a larger geographic scale as a means to limit the impacts of gene flow between populations. At the intermediate geographic scale, neither geographic origin of host, symbiont, nor an interaction between host and symbiont origin had a significant effect on host survival (Table 1; Figures 6, S7). We also observed no interaction between geographic origin of host and symbiont for host survival (Table 1; Figures 6, S7). Furthermore, the proportion of hosts surviving to adult did not differ between sympatric and allopatric combinations of host and symbiont (Table 1; Figure 6). For host survival to adult, we observed a significant effect of host geographic origin (χ^2^ = 12.89, df = 2, p = 0.002; Table 1, Figures 6, S7). However, we observed no effect of geographic origin of symbiont nor an interaction between the geographic origins of host and symbiont (Table 1; Figures 6, S7). Taken together, these results do not provide support for genetic specificity between host and symbiont lineages nor local adaptation at the intermediate geographic scale.

**Figure 6.**
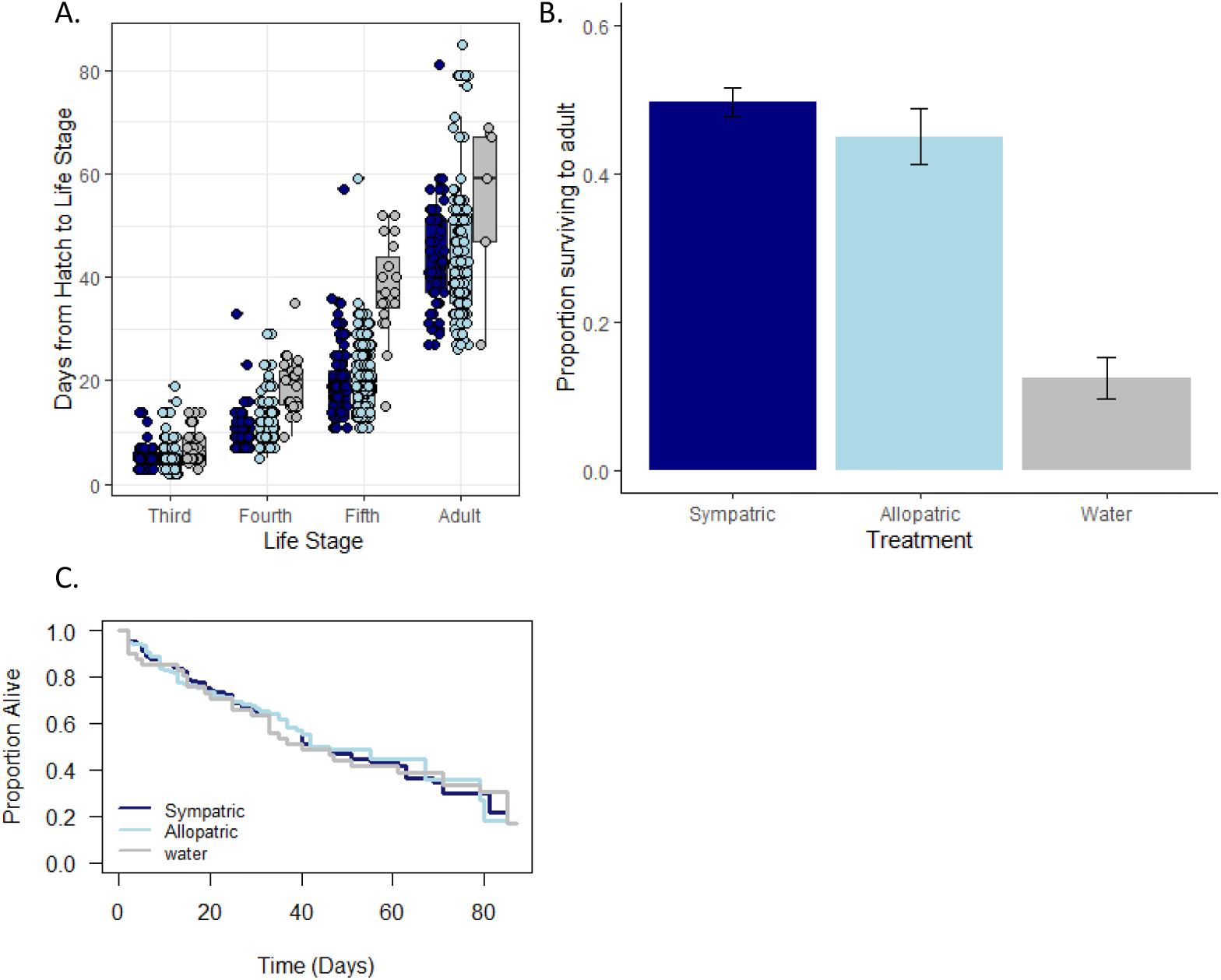
Host fitness at the intermediate geographic scale. Plot A shows overall host development rate when paired with sympatric versus allopatric symbionts (See figure S8 for all host development rate for all pairwise combinations of host and symbiont). We performed a cox proportional hazard model to determine whether host development rate to adult varied in response to host origin, symbiont origin, or an interaction between host and symbiont origin. We observed a significant effect of host geographic origin for rate of development to adult (p = 0.002). Bugs receiving water developed slower than those receiving a symbiont. Plot B shows the proportion of bugs surviving to adult across experimental treatments. We performed a cox proportional hazard model to determine whether host survival varied in response to host origin, symbiont origin, or an interaction between host and symbiont origin. Survival to adult for bugs receiving water was significantly reduced compared to those receiving a symbiont (p < 0.0001) but did not differ between sympatric and allopatric treatments. Survival over time did not vary across bugs receiving water versus those receiving a symbiont (C). This trend was driven by bugs that became developmentally “stuck” at a juvenile life stage but took a long time to die. We did not observe a significant effect of host origin, symbiont origin, nor an interaction between host and symbiont origin for host survival. We were unable to collect symbiont fitness data due to lab closures resulting from the COVID-19 pandemic.

#### Tests for local adaptation at the large geographic scale

Morphological variation exists between *A. tristis* squash bugs located in the eastern versus the western United States. Western bugs exhibit increased body size and melanization compared to eastern squash bugs. The phenotypic variation observed across Eastern and Western U.S. bugs suggests these populations are likely diverged from one another. We predicted phenotypic divergence across eastern and western host populations may result from divergent interactions with their microbial symbionts, so we repeated reciprocal inoculations at this largest geographic scale. We did not observe a significant effect of symbiont origin, host origin, nor an interaction between host and symbiont geographic origin for host survival (Table 1; Figure 7). Furthermore, there was no difference in the proportion of bugs that survived to adulthood between sympatric and allopatric combinations of host and symbiont (χ^2^ = 0.25013, df = 1, p = 0.617; Table 1; Figure 7). We observed a significant effect of host origin for development rate to adult (χ^2^ = 10.49, df = 1, p = 0.001; Table 1; Figure 7), such that hosts from the western United States developed slower than those from the eastern United States. We observed no effect of symbiont geographic origin nor a significant interaction between host and symbiont geographic origin for host development rate (Table 1; Figure 7). For symbiont fitness, we measured the number of CFUs per crypt of surviving adult squash bugs. We detected a significant effect of symbiont origin (F_1,15_ = 8.52, p = 0.01), but we did not observe a significant effect of host origin nor an interaction between host and symbiont. Overall, we did not find support for our prediction that pairwise coevolution underlies the maintenance of this horizontally transmitted mutualisms.

**Figure 7.**
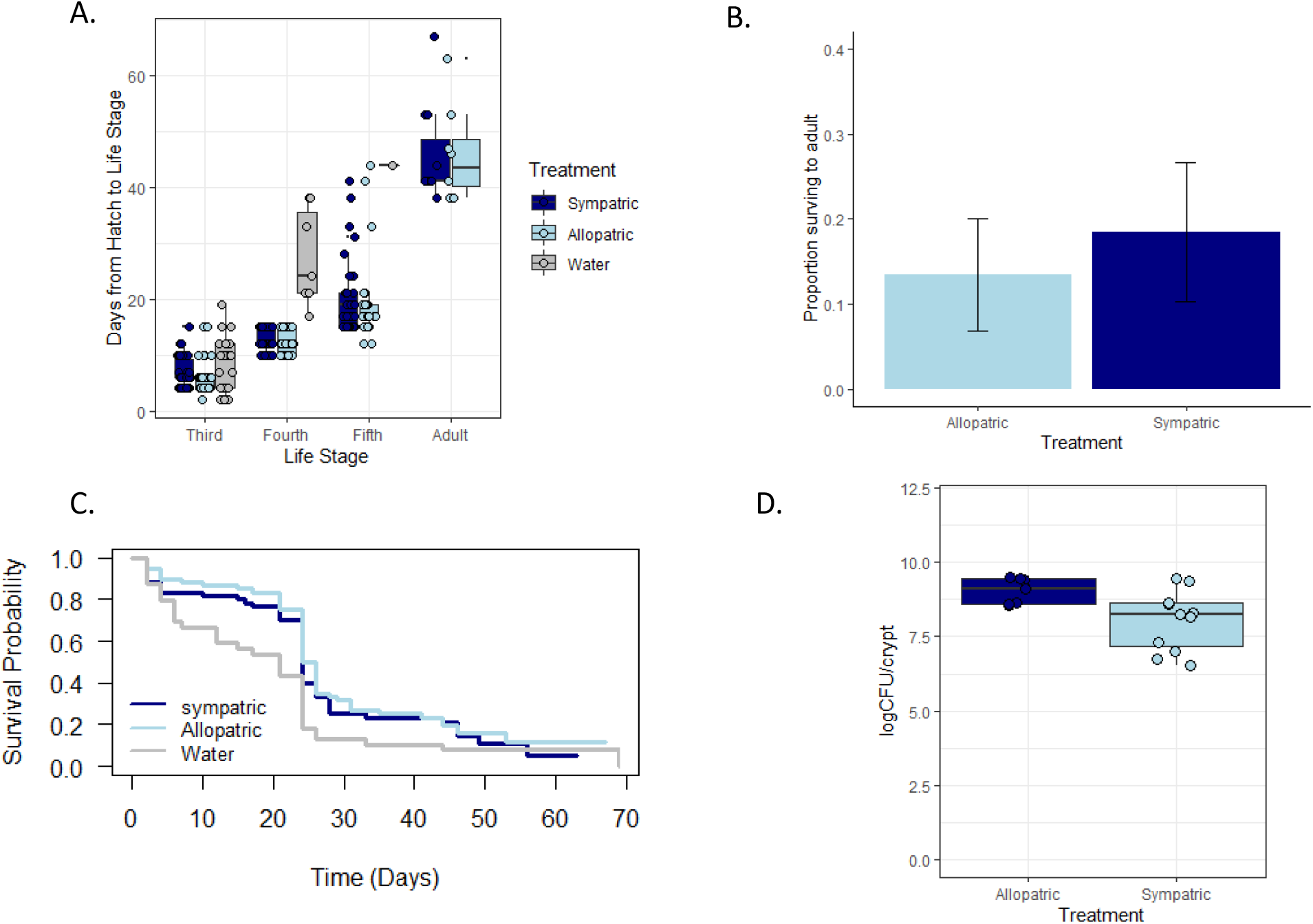
Host fitness at the large geographic scale. Plot A shows host development rate across all life stages when paired with a sympatric versus an allopatric symbiont. We performed a cox proportional hazard model to determine whether rate of development to adult varied in response to host origin, symbiont origin, or an interaction between host and symbiont origin. Plot B shows the proportion of bugs surviving to adult across experimental treatments. We detected no difference in survival to adult between sympatric and allopatric treatments (χ^2^ = 0.25, df = 1, p = 0.62). No bugs receiving water survived to adult, so they are not depicted here. Plot C shows survival across all experimental treatments. We performed a cox proportional hazard model to determine whether survival varied in response to host origin, symbiont origin, or an interaction between host and symbiont origin. We detected a significant effect of host origin (p = 0.001) on host fitness. Survival over time did not vary across bugs receiving water versus those receiving a symbiont. This trend was driven by bugs that became developmentally “stuck” at a juvenile life stage but took a long time to die. We did not observe a significant effect of host origin, symbiont origin, nor an interaction between host and symbiont origin for host survival. Plot D shows symbiont fitness for sympatric and allopatric treatments. We observed a significant effect of symbiont (F_1,15_ = 8.52, p = 0.01) on symbiont fitness but no effect of host nor an interaction between host and symbiont fitness.

### Specialization is not observed between host species and associated symbiont strains

We tested for specialization between symbionts and the host species from which they originated. If interactions were specialized, we predicted we would observe higher fitness interactions between hosts and conspecific-derived symbionts relative to symbionts derived from a heterospecific host. We did not detect a significant effect of symbiont origin for host survival across *A. tristis*, *A. andresii*, or *A. scorbutica* hosts (Table 2; Figures 8, S8). For all three host species, the proportion of hosts surviving to adult did not vary in response to receiving a symbiont from a conspecific host versus a heterospecific-derived symbiont (Figure 8). Similarly, symbiont origin did not affect host development rate to adult across all host species (Table 2; Figures 9, S8). Moreover, symbiont fitness (logCFU/crypt) did not vary in response to host species (*A. tristis*: F_2,17_ = 1.93, p = 0.18; *A. scorbutica*: F_2,17_ = 2.88, p = 0.08; *A. andresii*: F_2,35_ = 2.56, p = 0.09; Figures 9, S9). These results suggest hosts from each species mutualistically interact with a shared generalist symbiont.

**Figure 8.**
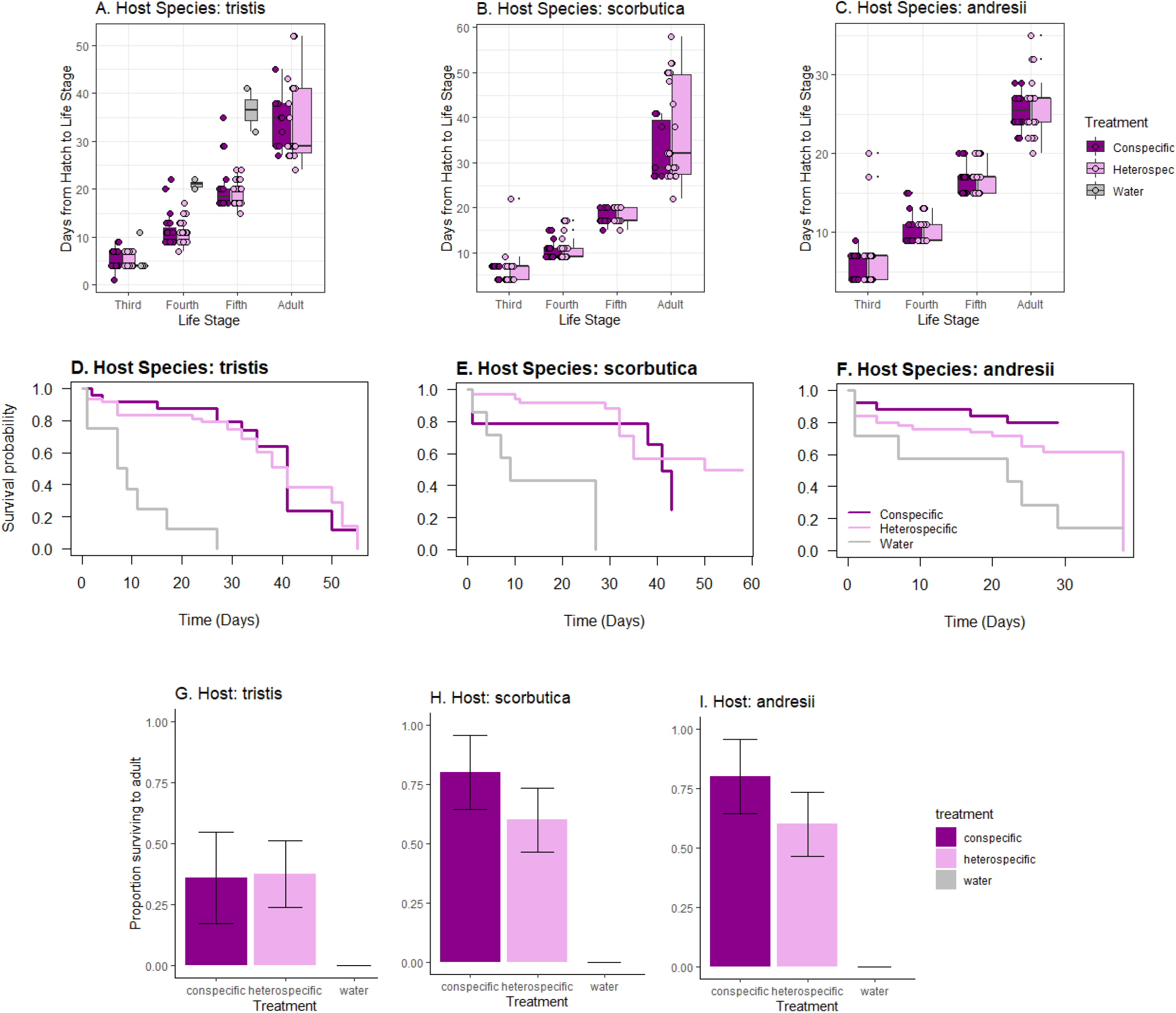
Host fitness data for reciprocal inoculations to test for specialization across host species with symbiont strain. Plots A-C show host development rate when paired with symbionts derived from a conspecific versus heterospecific host across all developmental stages for *A. tristis* (A), *A. scorbutica* (B), and *A. andresii* (C). We performed a cox proportional hazard model to determine whether symbiont origin had an effect on host rate of development to adult. We observed no effect of symbiont on host development rate to adult (see figure S8 for pairwise combinations of host species and symbiont strain). Plots D-F show host survival for *A. tristis* (A), *A. scorbutica* (B), and *A. andresii* (C) when paired with symbionts derived from a conspecific versus heterospecific host. We performed a cox proportional hazard model to determine whether host survival varied in response to symbiont origin. We observed no effect of symbiont origin on host survival (see figure S8 for pairwise combinations of host species and symbiont strain). Plots G-I show the proportion of bugs surviving to adult across all experimental treatments. We observed no difference in host survival when paired with symbionts isolated from a conspecific versus a heterospecific host.

**Figure 9.**
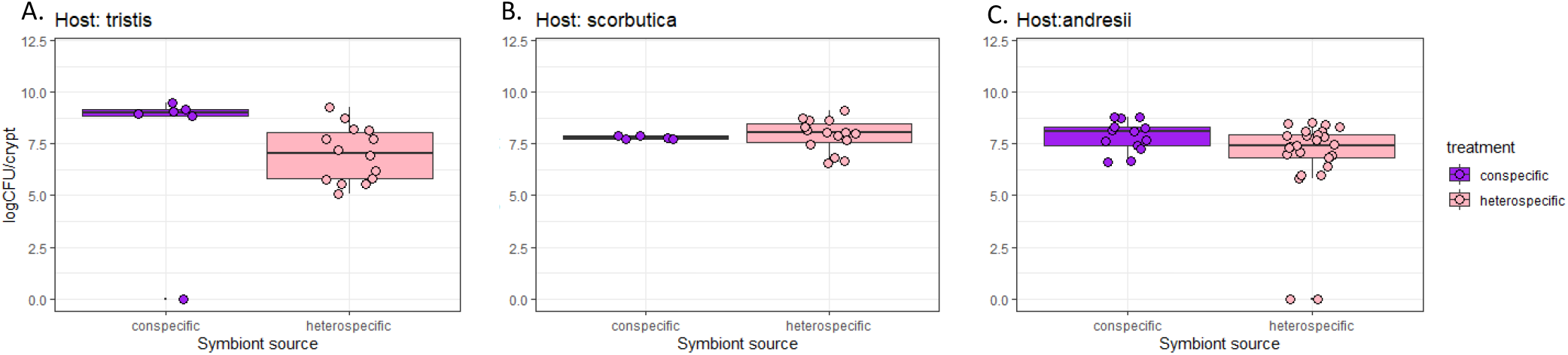
Symbiont fitness (logCFU/crypt) when paired with hosts that are conspecific or heterospecific to the hosts from which they were derived. Symbiont fitness was measured by dissecting the crypts of bugs that survived to adult during host fitness assays and counting the number of CFUs per crypt for each host species: *A. tristis* (A), *A. scorbutica* (B), *A. andresii* (C). We performed a quasipoisson distributed generalized linear model to determine whether symbiont fitness varied in response to host species. We observed no effect of host species on symbiont fitness (see xFigures S11 for all pairwise combinations).

## Discussion

In this study, we tested the hypothesis that coevolution underlies the maintenance of horizontally transmitted mutualisms. We assessed the role of pairwise coevolution by measuring host-symbiont specificity across three geographic scales. We observed no specificity between lineages. We then tested for specialization between three host species, *A. tristis*, *A. scorbutica*, and *A. andresii* with their associated *Caballeronia* symbionts but observed no evidence for specialization. Our results strongly demonstrate a lack of host-symbiont specificity in these interactions and fail to provide support for our hypothesis that pairwise coevolution underlies this horizontally transmitted mutualism. Instead, we observe evidence of generalist, beneficial symbionts likely under selection from a range of hosts for fixed phenotypic traits. These dynamics suggest the potential for diffuse coevolution across these interactions, such that the evolutionary trajectories of pairwise interactions are coupled through reciprocal selection between shared *Caballeronia* symbionts with each host species.

Horizontally transmitted symbiotic mutualisms are often characterized by genetic specificity, indicating a likely role for coevolution (Lee and Ruby 1994*b*; Aanen et al. 2002; Heath and Tiffin 2009; Murfin et al. 2015). Several studies involving plants and their microbial symbionts have directly tested for coevolutionary change between mutualistic hosts and symbionts by measuring geographic variation and local adaptation (Hoeksema and Thompson 2007; Barrett et al. 2012; Harrison et al. 2017*a*, 2017*b*; Rekret and Maherali 2019). Across these mutualistic interactions, evidence for local adaptation has been inconsistent, such that it is context-dependent (Rekret and Maherali 2019), one-sided (Hoeksema and Thompson 2007), or not observed (Barrett et al. 2012; Harrison et al. 2017*a*). In general, studies of local adaptation in horizontally transmitted mutualisms have provided little evidence for pairwise coevolution between hosts and symbionts.

In contrast, ample evidence for pairwise coevolution through tests of local adaptation has been observed across many host-parasite interactions (Ebert 1994; Thrall et al. 2002; Lively et al. 2004; Little et al. 2006; Greischar and Koskella 2007; King et al. 2009). The disparity in evidence for local adaptation within parasitic versus mutualistic interactions may result, in part, from the paucity of studies testing for local adaptation across mutualisms. Future work should continue to test mutualistic interactions for local adaptation to understand the coevolutionary dynamics underlying these interactions. Moreover, mutualistic and parasitic interactions are characterized by fundamentally different dynamics that may further account for this disparity. Parasites benefit from genetically matching their hosts, but hosts experience substantial fitness costs from matching their parasites. Interaction with parasites then selects for the maintenance of genetic variation in host populations to evade infection (Dybdahl and Lively 1996, 1998; Carius et al. 2001; King et al. 2009; Thrall et al. 2012). This asymmetric dynamic of costs and benefits between hosts and symbionts selects for the maintenance of phenotypic variation within and across populations (Yoder and Nuismer 2010). In mutualistic interactions, presumably the host and symbiont benefit from matching one another. Therefore, coevolution between hosts and symbionts is predicted to reduce phenotypic variation over time within and across mutualistic populations (Yoder and Nuismer 2010). The loss of phenotypic variation across mutualisms may result from interactions reaching their fitness optima. As a result, it may be difficult to observe coevolution in long-established mutualisms, despite coevolution potentially playing a prominent role in the initial establishment of the interaction. We observe similar patterns of genetic divergence for hosts and symbionts across populations at the intermediate geographic scale, but we do not observe correlated phenotypic divergence across these interactions. It is possible that phenotypic variation is not observed as previously coevolved traits involved in the mutualistic interaction are under stabilizing selection and genetic divergence across populations is driven by selection outside the interaction.

The success of mutualistic interactions is also often context-dependent (Thompson 1988; Bronstein 1994; Heath and Tiffin 2007; Hendry et al. 2014; Shantz and Burkepile 2014; Lowe et al. 2016). Genetic specificity across mutualisms may exist, but the phenotypic expression of differential fitness outcomes between specific host-symbiont combinations often depends on the external environment. For example, Rekret and Maherali (2019) tested for local adaptation between the plant host *Lobelia siphilitica* and its arbuscular mychorrhizal fungi symbionts. Local adaptation between host and symbiont was only observed at low phosphorous concentrations. Therefore, studies of local adaptation may depend on assessing the environmental context in which mutualists derive the greatest benefit from one another. We tested for local adaptation in a single common garden environment, possibly limiting some environmental component important for the expression of differential phenotypic traits across host and symbiont interactions. In general, evaluating the role of coevolution across symbiotic mutualisms may depend on using a broader approach.

Assessing the role of coevolution will likely depend on developing broad, new approaches that consider the unique dynamics of mutualisms. These empirical approaches must consider the effects of genetic specificity and environment in both newly established and ancient mutualisms to determine how mutualisms both establish and persist. The use of experimental evolution, as well as theoretical and empirical methods for assessing the evolutionary genetics dynamics of mutualism will be useful for assessing how coevolution underlies mutualisms (Brockhurst and Koskella 2013; Heath and Stinchcombe 2014; Hoang et al. 2016; Stoy et al. 2020). Moreover, a shortcoming of our tests for genetic specificity is the use of a single symbiont strain from each geographic location. By limiting reciprocal inoculations to a single strain, we may fail to capture phenotypic and genetic variation present within and across symbiont populations. We attempted to capture variation by picking genetically distant strains; however, because symbiont selection was dependent on variation in the 16s gene, it may not represent genetic variation relevant to the mutualistic interaction. In the future, our research will focus on characterizing genes underlying the interaction, variation at these loci, and assessing whether variation at specific loci corresponds to phenotypic variation across symbiont strains.

There has been a long recognized need to assess the coevolutionary dynamics between mutualists within a community context (Thompson 1994; Bronstein et al. 2003; Thrall et al. 2007). We tested for specialization between three *Anasa* host species with their respective *Caballeronia spp*. symbionts to determine the role of pairwise coevolution at the host genus level. Across our mutualistic interactions, we did not observe an effect of symbiont origin on host fitness, nor host species on symbiont fitness. Our results demonstrate that these interactions are not specialized. Rather, selection from multiple hosts likely selects for fixed cooperative traits in a shared generalist symbiont. Therefore, we found no evidence for pairwise coevolution. However, our results suggest the potential for diffuse coevolution between these hosts species with their common symbiont. Reciprocal selection between this range of insect hosts with their shared symbiont may maintain the interactions through diffuse coevolution.

Diffuse coevolution between multiple hosts and a shared symbiont can maintain overall benefits but strength of selection across pairwise interactions is reduced (Hougen-Eitzman and Rausher 1994; Iwao and Rausher 1997; Inouye and Stinchcombe 2011). As a result, phenotypic variation across interactions would not be observed. Therefore, diffuse coevolution may result in the maintenance of cooperative traits but limit opportunities for symbionts to become highly adapted and exploitative toward any single host. Specialization through pairwise coevolution may present significant costs for hosts dependent on environmentally acquiring symbionts in variable or unstable environments (Douglas 1998; Schwartz and Hoeksema 1998). As such, specialization may be especially costly for agricultural pests, such as squash bugs. Insects adapted to an agricultural setting must contend with sporadic availability of plant resources, variable crop variety, and enumerable pest mitigation methods. Diffuse coevolution with a generalist symbiont increases the probability that hosts will form successful mutualistic interactions each generation within these environments. Interacting with a variety of hosts may also increase the probability for symbiont transmission across patchily distributed agricultural plots.

We provide evidence of the potential for diffuse coevolution in the interaction between *Caballeronia spp*. and its insect hosts. However, we do not demonstrate direct evidence for diffuse coevolution. Future work should directly test the role of diffuse coevolution for the maintenance of horizontally transmitted mutualisms. This can be accomplished using experimental and evolutionary genetics approaches. Empirical methods rely on demonstrating that the strength of selection on the symbiont is altered by the presence or absence of a host species (Hougen-Eitzman and Rausher 1994; Iwao and Rausher 1997). Evolutionary genetics techniques can be used to determine whether the evolutionary trajectories of each pairwise symbiotic interaction are correlated due to shared underlying genetic interactions with the common partner (Hougen-Eitzman and Rausher 1994; Iwao and Rausher 1997; Inouye and Stinchcombe 2011; Ossler and Heath 2018). Moreover, as discussed previously, future work would benefit by assessing the effect of symbiont on host fitness using multiple strains from each host species, which may capture greater variation across symbiont strains.

The persistence of horizontally transmitted mutualisms presents a long-standing paradox. Theory has suggested that the maintenance of these interactions may result from coevolution across spatially structured populations of host and symbiont (Wilkinson 2001). We tested this hypothesis but found no evidence for host-symbiont specificity at the level of host species and host genus. Rather, our results suggest that interactions with a generalist symbiont produce the same fitness outcomes regardless of host or symbiont origin. Our results demonstrate a potential for diffuse coevolution, which we advocate should be the focus of future work aiming to elucidate the pathways by which horizontally transmitted mutualisms persist.

---

**Figure S1.**
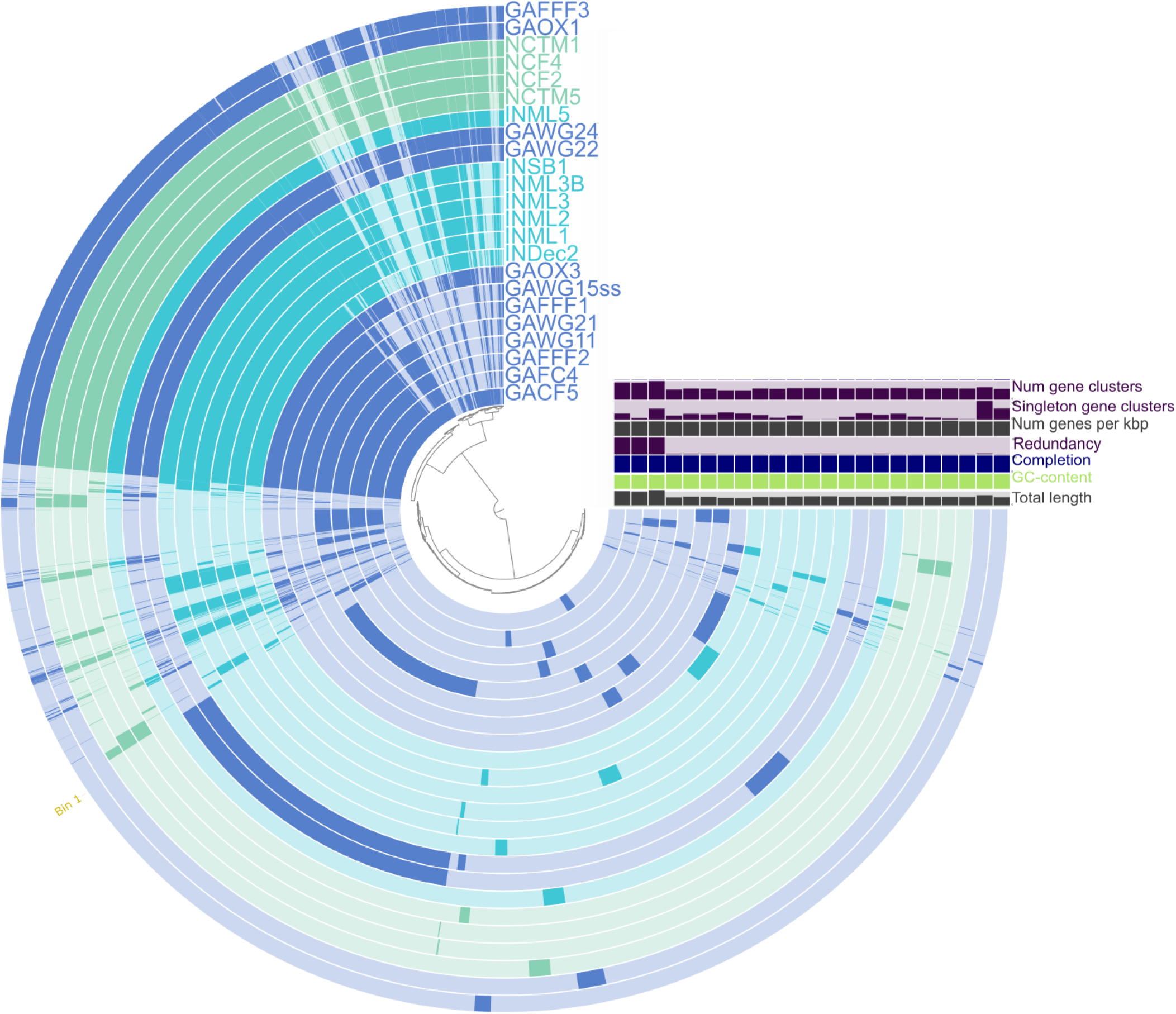
Pangenome analysis of symbiont strains isolated from IN (blue), GA (purple), and NC (green). Dark regions show the presence of gene clusters, and faded regions indicate the absence of gene clusters. The box to the right shows the relative number of gene clusters across genomes, relative number of genes present in only one genome (singleton genes), relative number of genes per kbp, redundancy, genome completion, relative GC-content, and total length of the sequence. Variation can be observed across the genomes of symbionts isolated from across their geographic range.

**Figure S2.**
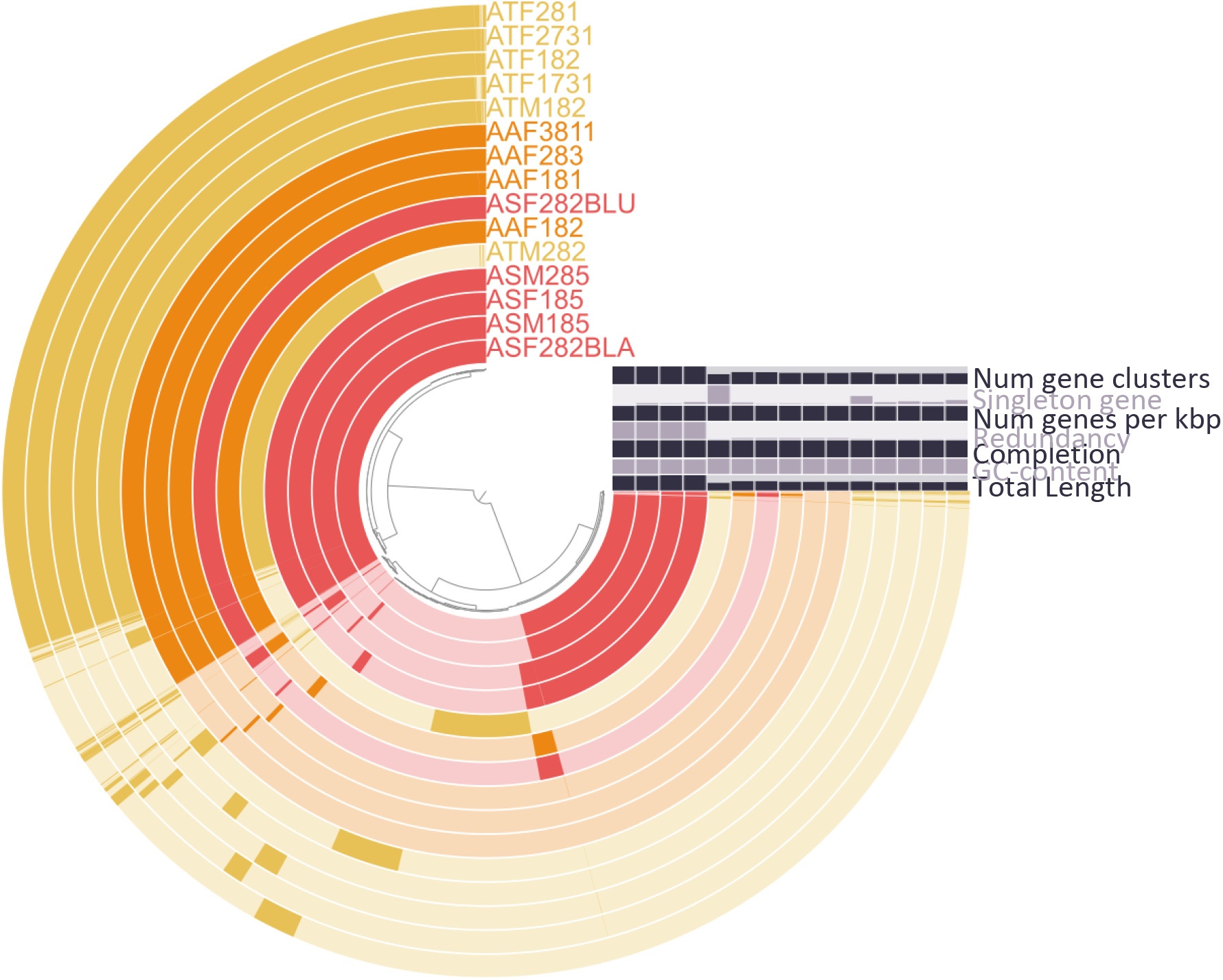
Pangenome analysis of symbiont strains isolated from three different host species *A. tristis* (AT; yellow), *A. andresii* (AA; orange), and *A. scorbutica* (AA; pink). Dark regions represent the presence of gene clusters and faded regions represent the absence of gene clusters. The box to the right shows the relative number of gene clusters across genomes, relative number of genes present in only one genome (singleton genes), relative number of genes per kbp, redundancy, genome completion, relative GC-content, and total length of the sequence. Variation can be observed across the genomes, particularly by the presence of gene clusters in *A. scorbutica* that are not present in *A. tristis* or *A. andresii* strains. Overall, little variation exists outside of these regions of the genome.

**Figure S3.**
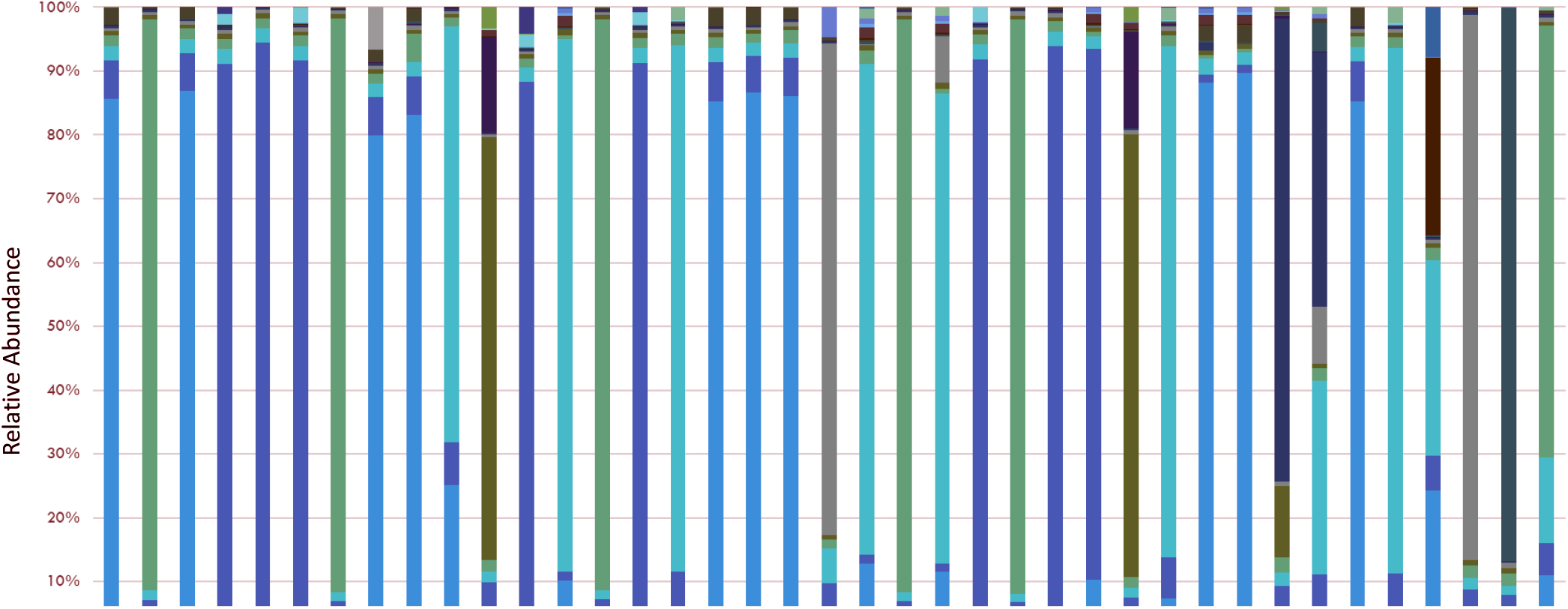
Relative abundance of OTUs within the *Burkholderiaceae* (Bacterial family containing *Caballeronia spp.*) across the crypts isolated from 39 *A. tristis* samples originating in Georgia (GA), Indiana (IN), and North Carolina (NC). Analysis was limited to the top twenty OTUs. Each bar represents the variation in *Burkholderiaceae* OTUs for a single sample. OTU analysis was performed using a sampling depth of 34,993 reads, which included 48% of all reads across 58 *Burkholderiacea* taxa. Analysis of the relative abundance of OTUs within the *Burkholderiaceae* across samples was randomized and limited to the top 20 OTUs, demonstrated here. The analysis indicates co-colonization of *Burkholderiacea* taxa commonly occurs within the crypts, despite observing little variation across cultivable strains. Moreover, this analysis indicates variation in crypt composition across individual bugs.

**Figure S4.**
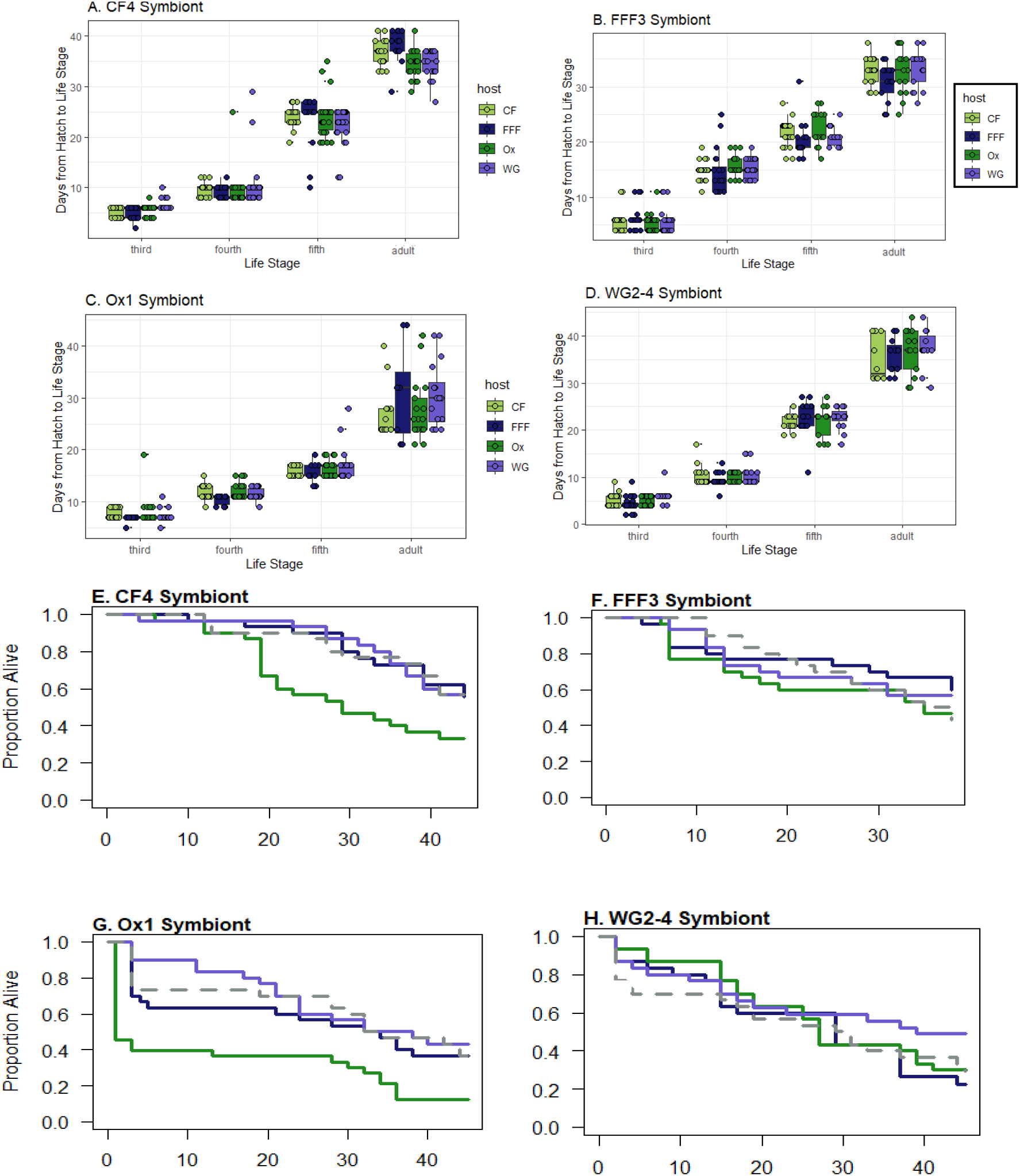
Host development rate and survival when paired with symbionts isolated from each site (CF, FFF, Ox, and WG) at the small geographic scale. Figures S4A-D show host development rate across all development stages. We performed a cox proportional hazard model to test for an effect of host origin, symbiont origin, and an interaction between host and symbiont origin on host development rate to adult. We observed a significant effect of symbiont origin (p < 0.001) and an interaction between host and symbiont origin (p = 0.03) for host development rate to adult. This interaction was not driven by differences in the effect of sympatric versus allopatric symbionts on host fitness (p = 0.23), as shown in Figures S4A-D. Figures S4E-H show host survival. We performed a cox proportion hazard mode to test for an effect of host origin, symbiont origin, and an interaction between host and symbiont origin on host survival. We observed a significant effect of symbiont origin (p = 0.0001) and host origin (p = 0.009) on host survival. Overall, we find no evidence for genetic specificity between host and symbiont.

**Figure S5.**
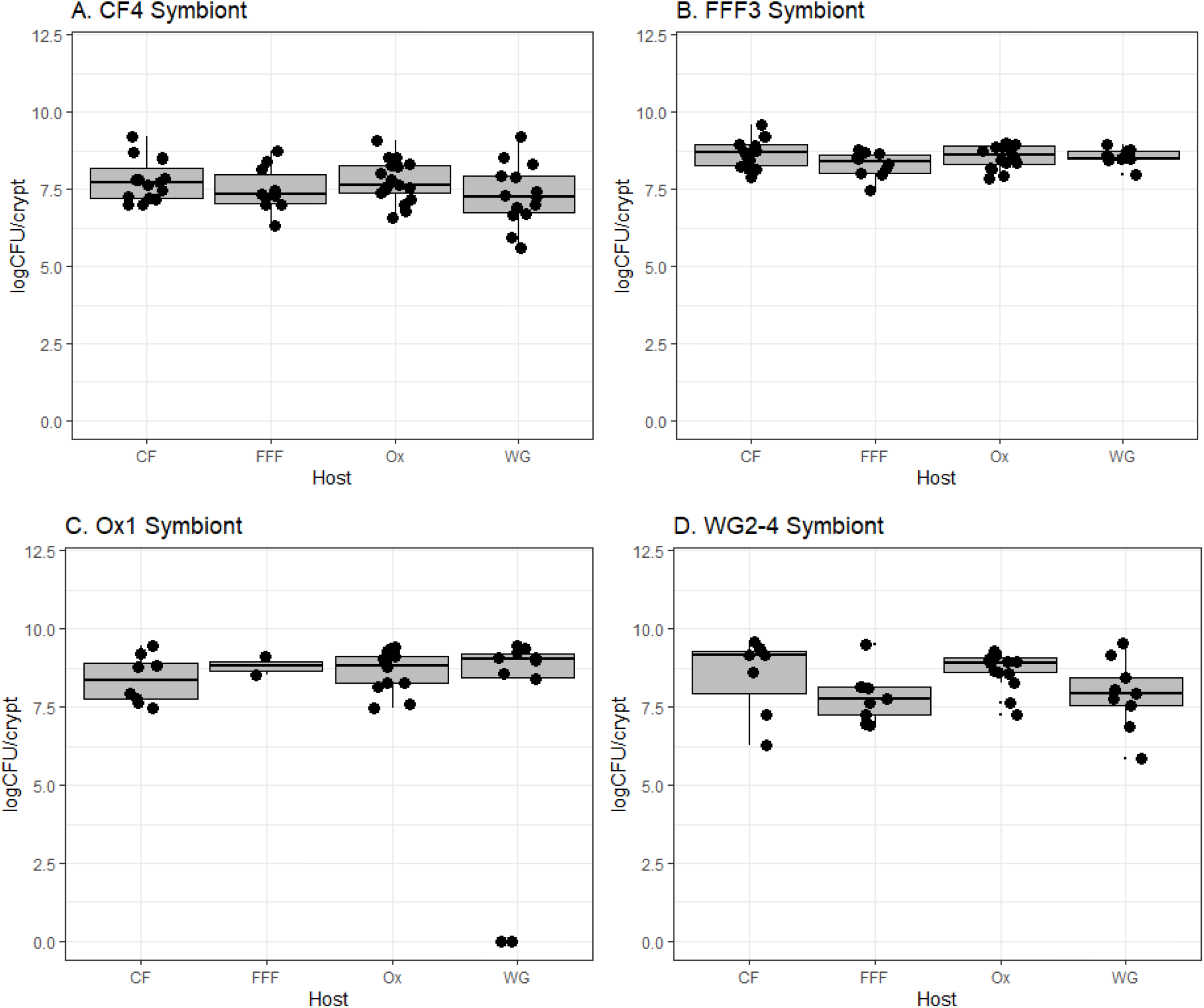
Symbiont fitness (logCFU/crypt) for the small geographic scale. We performed a quasipoisson distributed generalized linear model to determine whether symbiont fitness varied in response to host origin, symbiont origin, or an interaction between host and symbiont origin. Plots A-D demonstrate the fitness for each individual symbiont when paired with each host. Symbiont titer within crypts was measured by dissecting adult squash bugs. Symbiont fitness did not vary in response to host origin, providing no evidence for local adaptation.

**Table S1.** Reciprocal inoculation to test the effect symbiont on host survival and development rate at the small geographic scale when inoculations were performed synchronously. We performed a linear contrast to test whether interactions resulted from an effect of sympatric versus allopatric symbionts on host fitness.

| Test | Effect | df | $\chi^2$ | p-value |
| --- | --- | --- | --- | --- |
| Survival | Symbiont | 3 | 3.40 | 0.33 |
|  | Host | 1 | 0.03 | 0.86 |
|  | Host*Symbiont | 3 | 8.92 | **0.03\*** |
| *sympatric-allopatric contrast (p = 0.90) |  |  |  |  |
| Development Rate | Symbiont | 3 | 5.60 | 0.13 |
|  | Host | 1 | 2.58 | 0.12 |
|  | Host*Symbiont | 3 | 2.44 | 0.49 |

**Figure S6.**
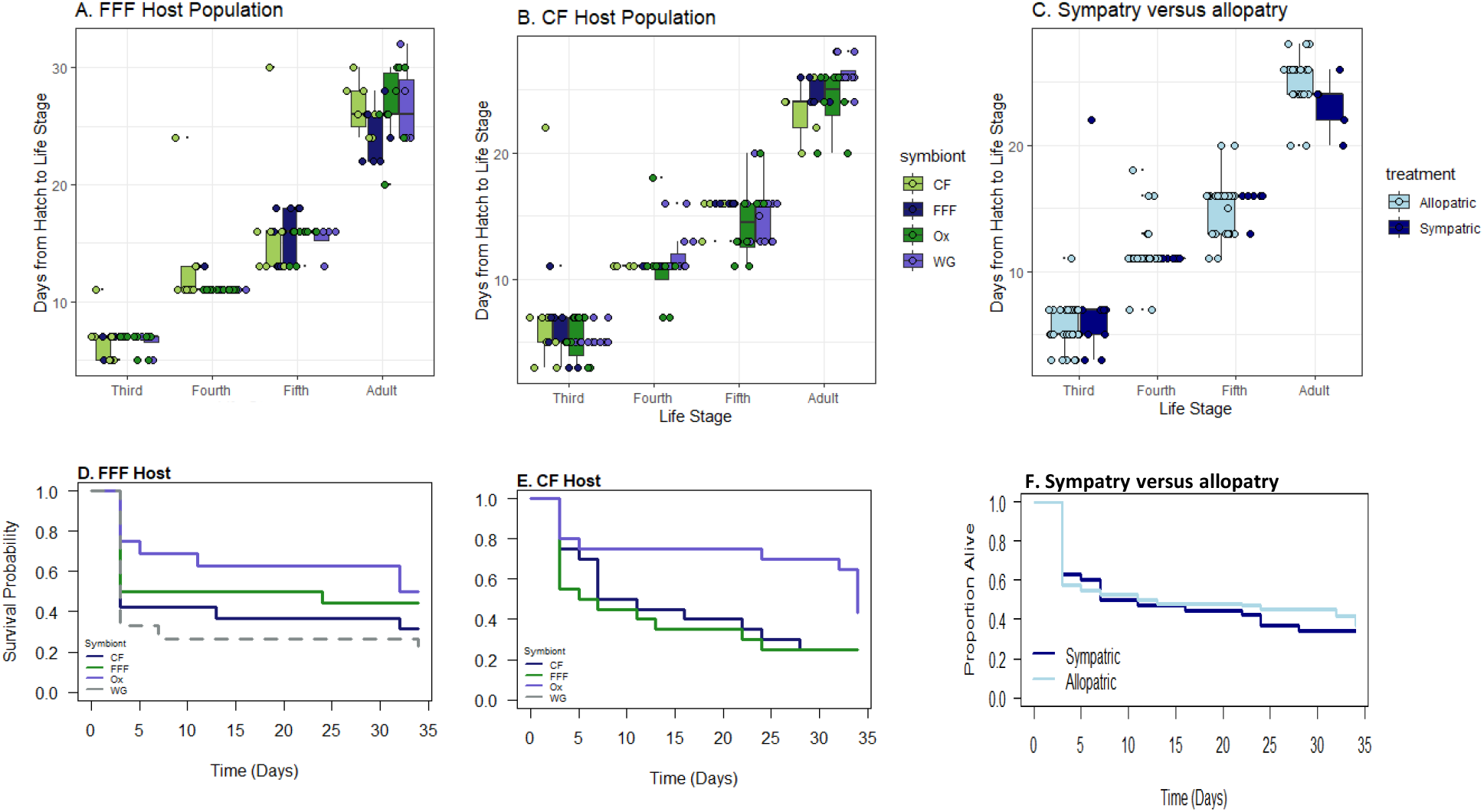
Host fitness at the small geographic scale for the FFF and CF host populations when inoculations with symbionts from each population were performed synchronously. Plots A and B show host development rate for the FFF and CF populations. We performed a cox proportion hazard model to determine whether host rate of development to adult varied in response to host origin, symbiont origin, or an interaction between host and symbiont origin. Development rate did not vary in response to the origin of host, symbiont, nor an interaction between the origin of host and symbiont. Plot C shows overall development rates for sympatric versus allopatric combinations of host and symbiont. Plots D and E show host survival for the FFF and CF populations. We performed a cox proportional hazard model to determine whether host survival varied in response to host origin, symbiont origin, or an interaction between host and symbiont origin. We observed no effect of host nor symbiont origin on host survival. However, we did observe an interaction between host and symbiont origin (p = 0.03), but this was not driven by differences in the effect of sympatric versus allopatric symbionts on host fitness (p = 0.90). Plot F shows the overall survival for sympatric versus allopatric combinations of host and symbiont.

**Figure S7.**
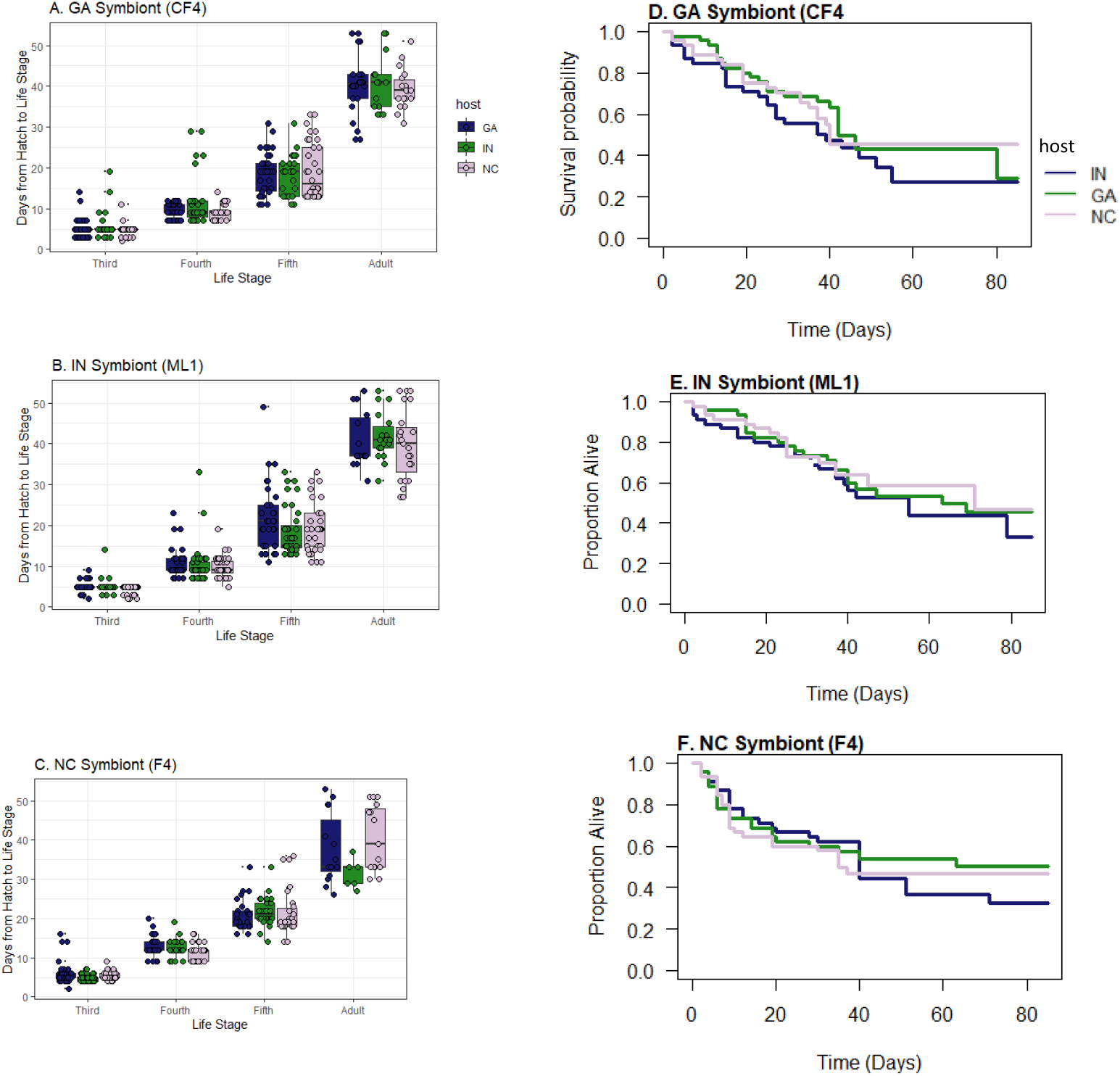
Host development rate and survival for all pairwise combinations of hosts and symbionts isolated across the three states (GA, IN, and NC) at the intermediate geographic scale. Plots A-C show host development rate across all developmental life stages. We performed a cox proportional hazard model to test for an effect of host origin, symbiont origin, or an interaction between host and symbiont origin for rate of development to adult. We detected a significant effect of host geographic origin on host development rate. We did not observe an effect of symbiont nor an interaction between host and symbiont geographic origin. Plots D-F show host survival. We performed a cox proportional hazard model to determine whether survival varied in response to host origin, symbiont origin, or an interaction between host and symbiont origin. Host survival did not vary in response to geographic origin of host, symbiont, nor an interaction between host and symbiont geographic origin.

**Figure S8.**
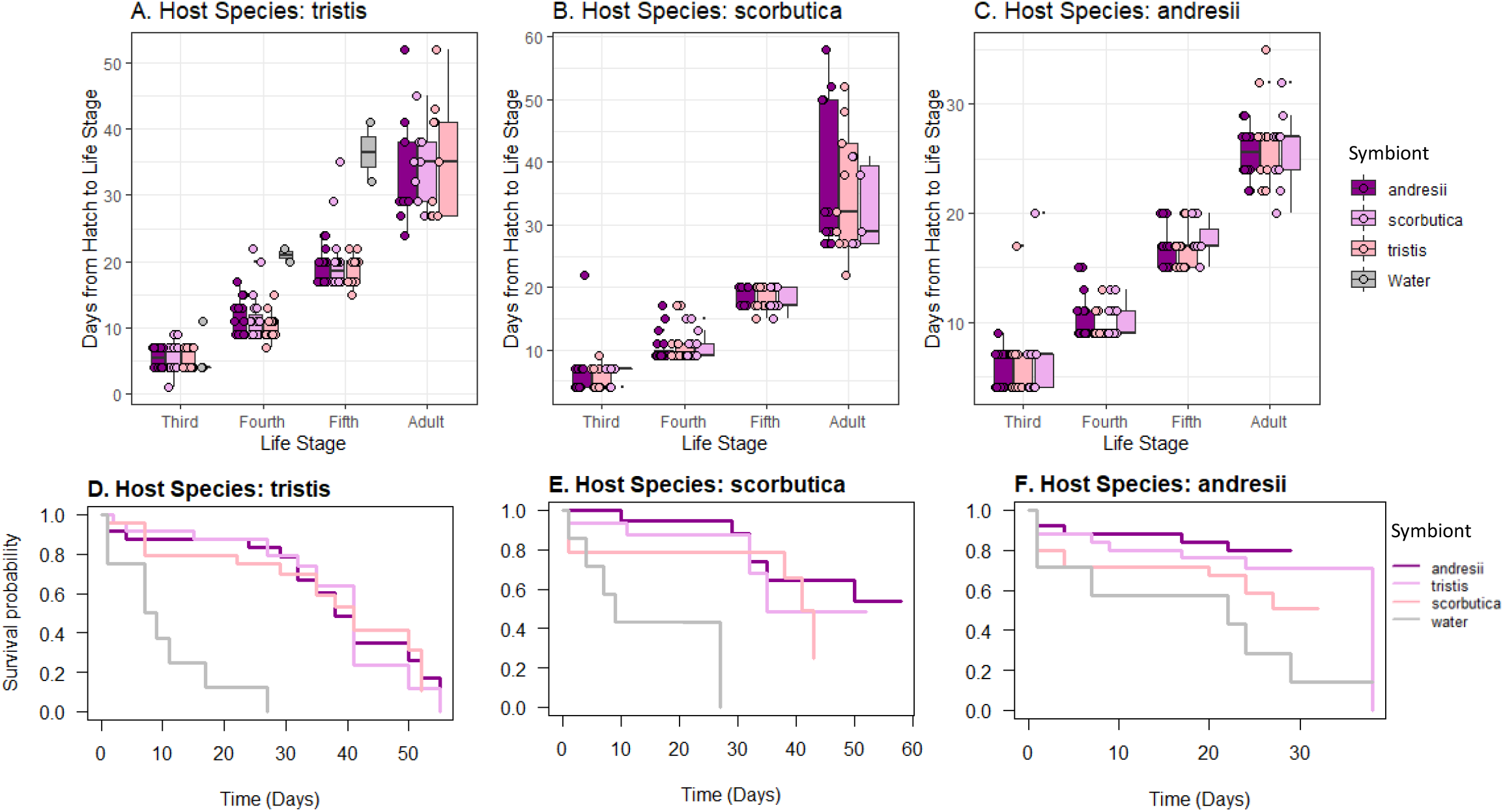
Host fitness data for all pairwise combinations of host species and symbiont strains isolated from heterospecific or conspecific hosts. Plots A-C show host development rate across all developmental life stages: *A. tristis* (A), *A. scorbutica* (B), and *A. andresii* (C). We performed a cox proportional hazard model to assess the rate of development to adult across treatments. We did not observe an effect of symbiont origin on host development rate. Plots D-F show host survival for each host species: *A. tristis* (D), *A. scorbutica* (E), and *A. andresii* (F). We performed a cox proportional hazard model to assess the effect of symbiont origin on host survival, and we did not observe a significant effect of symbiont origin on survival for any host species.

**Figure S9.**
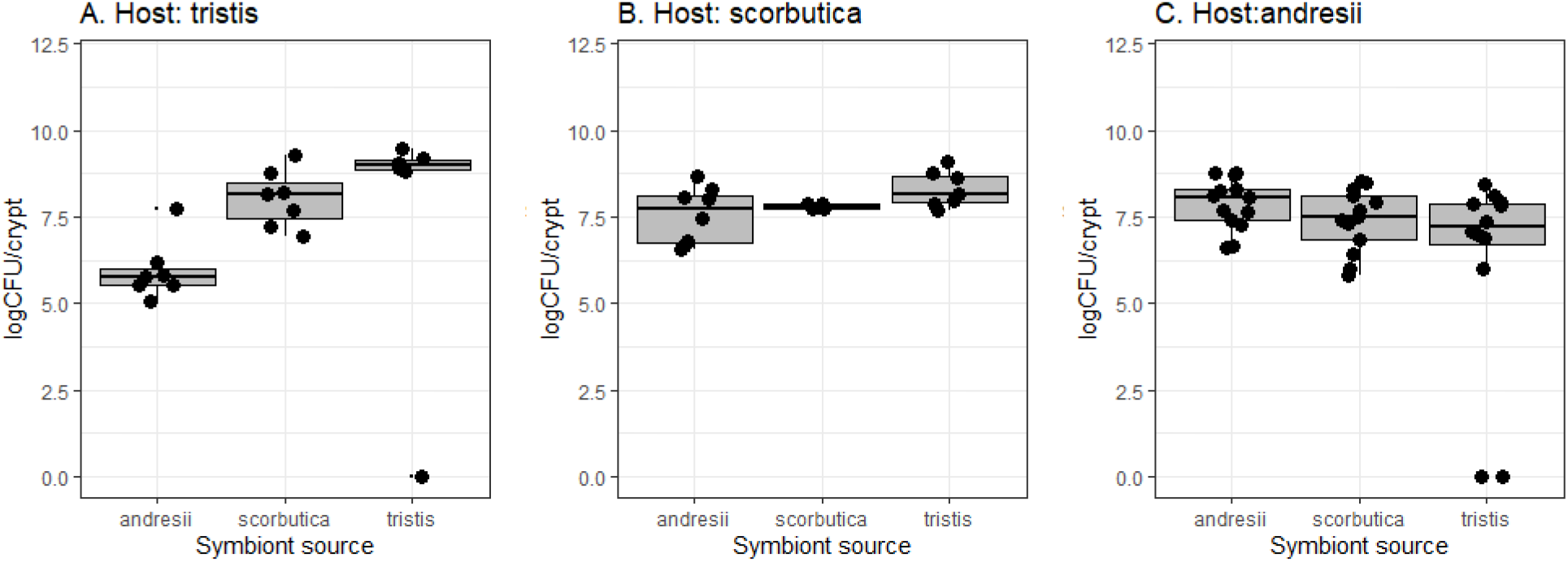
Symbiont fitness (logCFU/crypt) for all pairwise combinations of host species and symbiont origin. Crypts of squash bugs that survived to adult were dissected, crushed, and plated to assess the number of symbiont CFUs per crypt. The fitness of symbionts isolated from each host species is shown when symbionts were paired with *A. tristis* (A), *A. scorbutica* (B), and *A. andresii* (C). We performed a quasipoisson distributed generalized linear model to determine whether symbiont fitness varied in response to host species. We observed no effect of host species on symbiont fitness, regardless of the species from which the symbiont was isolated.

